# Osteochondroprogenitor cells and neutrophils expressing p21 and senescence markers modulate fracture repair

**DOI:** 10.1101/2024.02.01.578420

**Authors:** Dominik Saul, Madison L. Doolittle, Jennifer L. Rowsey, Mitchell N. Froemming, Robyn L. Kosinsky, Stephanie J. Vos, Ming Ruan, Nathan LeBrasseur, Abhishek Chandra, Robert Pignolo, João F. Passos, Joshua N. Farr, David G. Monroe, Sundeep Khosla

## Abstract

Cells expressing features of senescence, including upregulation of p21 and p16, appear transiently following tissue injury, yet the properties of these cells or how they contrast with age-induced senescent cells remains unclear. Here, we used skeletal injury as a model and identified the rapid appearance following fracture of p21+ cells expressing senescence markers, mainly as osteochondroprogenitors (OCHs) and neutrophils. Targeted genetic clearance of p21+ cells suppressed senescence-associated signatures within the fracture callus and accelerated fracture healing. By contrast, p21+ cell clearance did not alter bone loss due to aging; conversely, p16+ cell clearance, known to alleviate skeletal aging, did not affect fracture healing. Following fracture, p21+ neutrophils were enriched in signaling pathways known to induce paracrine stromal senescence, while p21+ OCHs were highly enriched in senescence-associated secretory phenotype factors known to impair bone formation. Further analysis revealed an injury-specific stem cell-like OCH subset that was p21+ and highly inflammatory, with a similar inflammatory mesenchymal population (fibro-adipogenic progenitors) evident following muscle injury. Thus, intercommunicating senescent-like neutrophils and mesenchymal progenitor cells are key regulators of tissue repair in bone and potentially across tissues. Moreover, our findings establish contextual roles of p21+ *vs* p16+ senescent/senescent-like cells that may be leveraged for therapeutic opportunities.

## INTRODUCTION

At the cellular level, aging is characterized by several hallmarks, including cellular senescence (1), which is driven by an increase of the cyclin-dependent kinase inhibitors (CDKIs) *Cdkn1a* (p21) and/or *Cdkn*2a (p16) (2). Senescent cells also demonstrate a senescence-associated secretory phenotype (SASP), consisting of pro-inflammatory cytokines, multiple chemokines, and matrix degrading proteins, driving tissue dysfunction in a paracrine and systemic manner (3). There is now considerable evidence that senescent cells accumulate with age across tissues and that clearance of these cells in mice using either genetic or pharmacologic approaches ameliorates multiple aging phenotypes, including frailty, osteoporosis, cardiovascular disease, metabolic dysfunction, and others (for a review, see DiMicco et al. (4)). This has led to intense interest in the development of compounds (“senolytics”) that target senescent cells to ameliorate these age-associated morbidities (5).

In addition to their role in aging, however, cells expressing features of senescence also appear rapidly following tissue injury and modulate the repair process (6). In contrast to age-associated senescent cells, these injury-related senescent-like cells remain relatively poorly characterized. Moreover, they appear to play variable roles in tissue repair, with evidence of a beneficial role in the healing of skin wounds (7) and lung repair following some (8) but not other forms (9) of lung injury *vs* a detrimental role in bone (10, 11) or muscle repair (12). In addition to understanding their fundamental biology, it is clearly important to characterize these injury-related senescent-like cells so we can better define the benefits *vs* risks of senolytic therapies for various age-associated morbidities as they rapidly move to the clinic.

Our group previously demonstrated (10) that cells expressing features of senescence, (*i.e.,* increased p16 and p21 expression, production of a SASP, and evidence of irreversible telomeric DNA damage [telomere-associated foci (TAFs) (13), arguably one of the definitive assays for senescent cells]) accumulate transiently in the fracture callus. Pharmacologically targeting these cells by senolytic drugs (dasatinib + quercetin [D+Q]) accelerated both the time-course and ultimate biomechanical strength of the healed fracture (10), as also confirmed by Liu et al. (11). In recent studies (14), we used cytometry by time-of-flight (CyTOF) as well as single cell RNA-sequencing (scRNA-seq) to rigorously define senescent cells in the context of aging at the single-cell level consistent with criteria outlined by the International Cell Senescence Association (15): upregulation of p16 and/or p21, growth arrest, upregulation of a SASP and anti-apoptotic pathways, and evidence of DNA damage. In the present study, we used these validated tools to define the cellular identity and functional characteristics of cells expressing features of senescence following tissue injury using fracture as our model and contrasted these injury-related senescent-like cells to senescent cells associated with aging. We first characterized cells expressing senescence markers, including p21 and p16, in the fracture callus using CyTOF. Next, given the increasing evidence that p21 and p16 may be different functionally (16), we compared the effects of genetic clearance of p21+ vs. p16+ cells on fracture healing using newly developed *p21-ATTAC* (17) as well as established *p16-INK-ATTAC* mouse models (18, 19). We further used CyTOF complemented by scRNA-seq analysis to provide a detailed characterization of the cells modulating fracture healing. Finally, we compared our findings following bone injury to available data in the context of muscle injury and identified very analogous mesenchymal progenitor populations in muscle that develop features of injury-related senescence, indicating that our findings likely extend across tissues. Collectively, our studies identify novel mesenchymal progenitor and immune populations expressing features of senescence following tissue injury that differ in important ways from classical senescent cells associated with aging, but which could nonetheless be targeted to enhance fracture healing and potentially facilitate repair in other tissues.

## RESULTS

### Appearance of p21+ and p16+ cells expressing features of senescence during fracture healing

To investigate potential injury-related senescent-like cells that arise during fracture healing, we performed CyTOF on single-cell suspensions from digested callus samples throughout the fracture healing process using antibodies, including for p16 and p21, that we have previously extensively validated (14). To capture the four stages of fracture healing (inflammatory, soft callus, hard callus, and remodeling phase), we performed a diaphyseal tibial fracture in 47 *C57BL/6* mice and harvested the newly formed callus on days 3, 7, 14 and 28. The minced and digested callus cells were analyzed by CyTOF (Figure 1A, Supplementary Figure 1A). We identified both mesenchymal and immune cell populations with unique abundance patterns throughout fracture healing (Figure 1A, B). p16 and p21, previously observed to be expressed in the fracture callus (10, 11) were expressed in both mesenchymal and immune cell populations, with osteochondroprogenitor (OCH) cells, osteoblasts, and monocytes/macrophages expressing high levels of p16, and OCH cells and neutrophils expressing high levels of p21 (Figure 1C, Supplementary Figure 1B). Importantly, we found that co-expression of SASP factors with p16 and p21 was particularly strong in OCH cells (Figure 1C), which highly expressed osteogenic (Runx2, Osterix, ALPL, DMP1), chondrogenic (Sox9, Sox6), and progenitor (CD200, PDGFRα, CD73) markers (Figure 1B).

**Figure 1:**
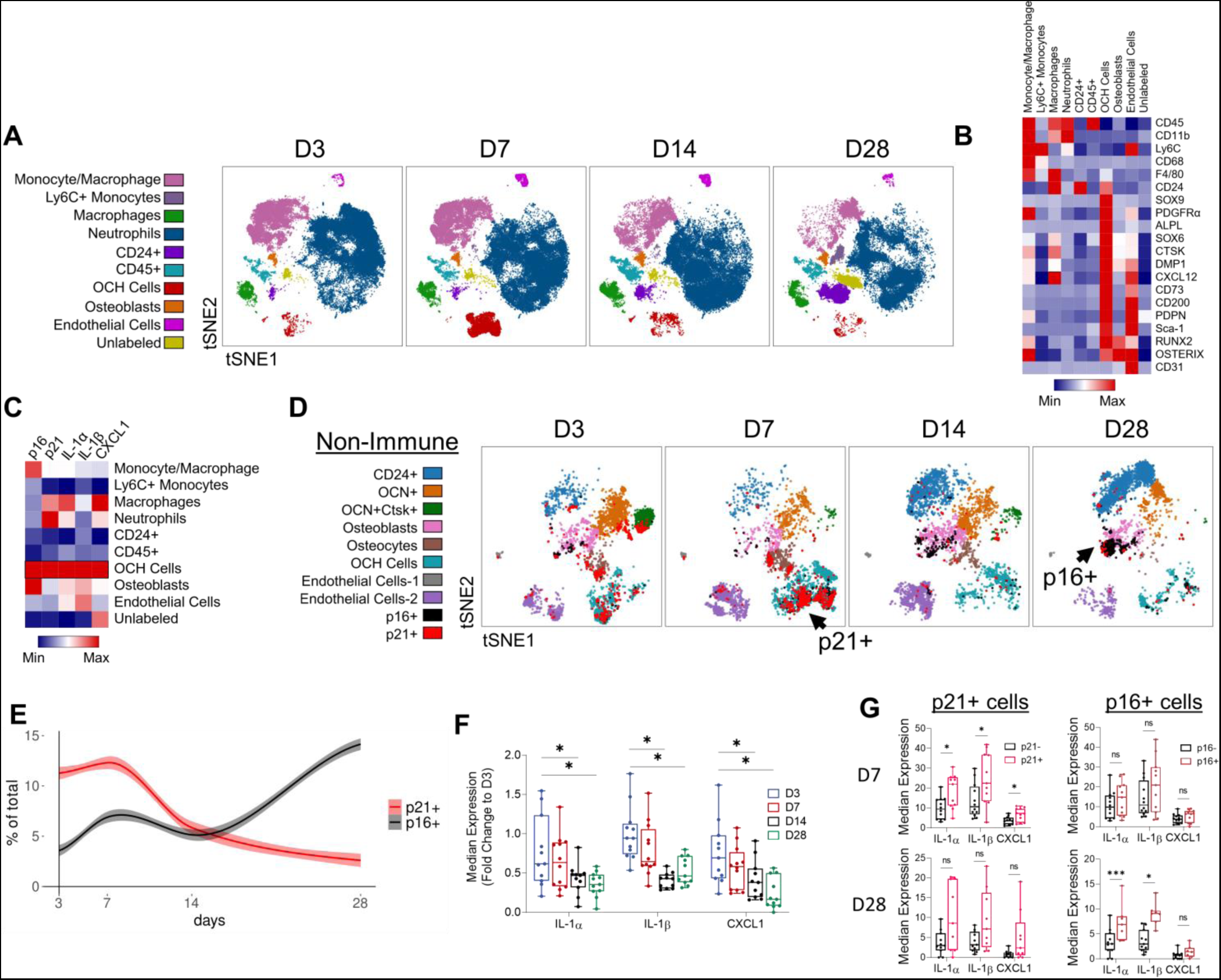
p21+ and p16+ cells appear in a divergent manner during fracture healing. (A) tSNE visualization of clustered cell populations across murine fracture healing by CyTOF. (B) Heatmap representation of identification and (C) senescence and SASP marker median expression across all clusters. (D) tSNE visualization of non-immune (CD45-CD11b-) callus cells across fracture healing, overlaid with p16+ (black) and p21+ cells (red). (E) p16+ and p21+ cell abundances across fracture healing in non-immune cells. (F) SASP marker median expression throughout fracture healing in all non-immune cells and (G) in p16+ and p21+ non-immune cells. Day 3: n=12 mice (6 female, 6 male), day 7: n=12 mice (6 female, 6 male), day 14: n=12 mice (6 female, 6 male), day 28: n=11 mice (6 female, 5 male). (F) One-way ANOVA with Tukey correction for multiple comparisons, (G) Mann-Whitney test. *p<0.05, ***p<0.001.

To investigate the mesenchymal progenitor cells at higher resolution, non-immune (CD45-CD11b-) callus cells were re-clustered and phenotyped for their senescence profile (Figure 1D, Supplementary Figure 1C-E). p21+ cells appeared early in fracture healing (D3-D7), predominantly as OCH cells, while p16+ cells appeared late (D14-28) as osteoblasts (Figure 1D). This temporal expression pattern for p16 and p21 was observed among all non-immune cells (Figure 1E). Notably, the existence of p21+ cells coincided with high inflammation, demonstrated by an early peak (D3) in expression of SASP proteins IL-1α, IL-1β, and CXCL1 (Figure 1F). Moreover, SASP markers were enriched in p21+ cells at an early stage (D3-7), but not at a late stage (D28), while the opposite was true for p16+ cells (Figure 1G).

### Genetic clearance of p21+, but not p16+, cells accelerates fracture healing

To evaluate the functional contribution of inflammatory p21+ versus p16+ cells toward fracture healing, we first leveraged the *p21-ATTAC* mouse model, recently validated by our laboratory (17) (Figure 2A). This *p21-ATTAC* mouse (analogous to the *p16-INK-ATTAC* mouse (18)) contains a “suicide” transgene driven by the *p21^cip1^* promoter, whereby administration of AP20187 (AP) induces caspase 8-driven apoptosis in p21+ cells. Figure 2A also shows the design of the study, including the time-points of twice-weekly AP administration following fracture. qRT-PCR analysis of the callus area demonstrated a marked reduction of both p21 mRNA (*Cdkn1a*) and the *p21-ATTAC* transgene (*eGFP*) following AP treatment (Figure 2B). In the five-week healing course of a transverse tibial fracture, we found that the x-ray-based healing score (20) was consistently improved in mice cleared of p21+ cells (Figure 2C). We next performed in-depth callus size measurements on a weekly basis and found the relative callus area to be significantly increased in the AP-treated mice from the second week onward (Figure 2D). At the conclusion of fracture healing, both callus bone volume (by µCT) and biomechanical stiffness were increased in mice cleared of p21+ cells (Figure 2E, F). Using fluorescent labeling of the newly formed callus area (21), we found an increase in the bone formation rate/bone surface (BFR/BS) and mineral apposition rate (MAR) in weeks 3 and 4 with clearance of p21^+^ cells (Figure 2G-I). Interestingly, we found no difference in osteoblast numbers between the treatment groups (Figure 2J), suggesting that the increases in MAR and BFR in AP-treated mice were due primarily to an increase in osteoblast activity, rather than number. In addition, the number of osteoclasts per bone perimeter were reduced in the AP-treated mice (Figure 2K).

**Figure 2:**
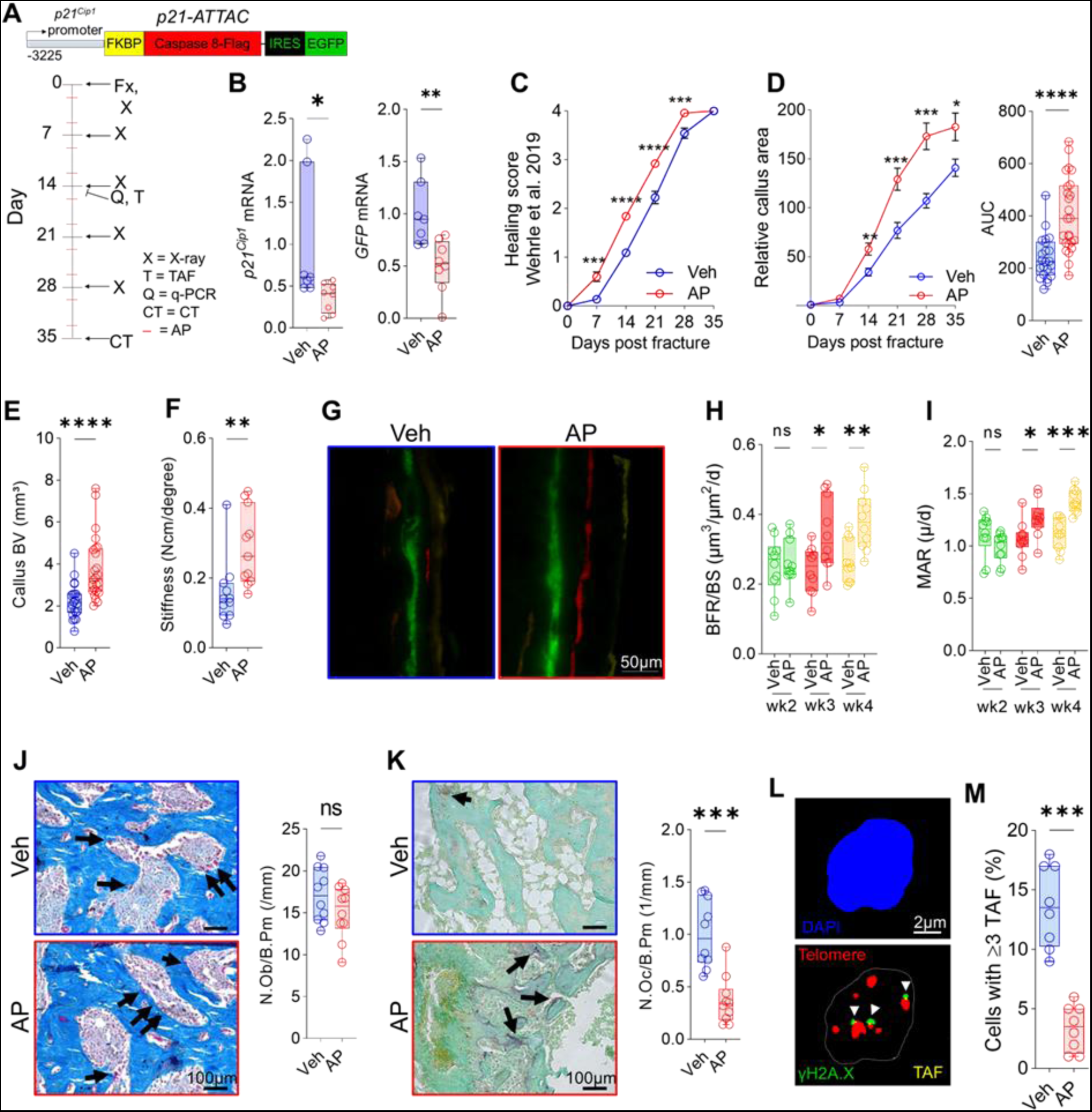
Clearance of p21+ cells accelerates fracture healing by increasing bone formation rates and reducing osteoclast numbers. (A) Schematic of the *p21-ATTAC* transgene and overall study design. 4-6 month old *p21-ATTAC* mice were used to selectively clear p21+ cells through AP administration twice weekly over a 5-week fracture healing timecourse. (B) qPCR measurement of *p21^Cip1^* and GFP (*p21-ATTAC* transgene) mRNA expression after AP treatment on day 14. (C) Fracture healing score (described by Wehrle et al. (20)), and (D) callus area as measured by weekly x-ray. (E) Micro-CT of callus bone volume. (F) Tibial stiffness measured by biomechanical testing. (G-I) Histomorphometric analysis of bone formation rate per bone surface (BFR/BS) and mineral apposition rate (MAR) through weekly injections of bone-labeling dyes (see Methods). (J) Histological quantification of osteoblasts through Masson trichrome staining. (K) Histological quantification of osteoclasts through tartrate-resistant acid phosphatase (TRAP) staining. (L) Telomere-associated foci (TAF) staining (D14) for DNA damage. (M) Quantification of cells exhibiting ≥3 TAF per cell. n=8-11 (B,F-M) or n=22-25 (C,D) per treatment, equally split by sex. (B,D,E,J-M) Mann-Whitney, (C,D) Two-way ANOVA + Sidak correction, (H,I) Multiple t-test + FDR correction. *p<0.05, **p<0.01, ***p<0.001, ****p<0.0001.

We next evaluated whether clearance of p21+ cells was associated with a reduction in senescent cell signatures. To do so, we performed an analysis of telomere-associated foci (TAF, Figure 2L), which identifies sites of irreversible telomeric DNA damage, a hallmark of senescent cells (13). We have previously demonstrated that TAFs increase markedly in the fracture callus, peaking at day 14 following fracture and subsequently returning towards baseline levels (10). Here, we observed a marked reduction of TAF+ cells in the AP- *vs.* the vehicle-treated mice at day 14 (Figure 2M). Thus, clearance of p21+ cells effectively reduced hallmarks of senescence, resulting in accelerated fracture healing and stronger bone.

To evaluate whether the clearance of p16+ cells also altered the time course of fracture healing, we used the *p16-INK-ATTAC* model (Supplementary Figure 2A). We confirmed adequate functionality of the *p16-INK-ATTAC* model by demonstrating downregulation of the *Casp8* portion of the transgene cassette in the fracture callus following AP treatment, consistent with clearance of cells expressing the *p16*-driven suicide transgene (Supplementary Figure 2B). Note that the primers used for *Casp8* are specific for the human transcript encoded by the transgene (22). Weekly x-ray analyses showed a small, but insignificant difference in the callus formation rate with a marginal acceleration in radiographic callus formation in the AP-treated group in the late stage (day 28, Supplementary Figure 2C). However, the resulting total callus volume of the healed bones remained unchanged (Supplementary Figure 2D). Both our CyTOF analyses and previous studies (10) demonstrated that p16+ cells begin to emerge within the callus around day 14 post-fracture. To detect the expected effect of AP on senescent cells, we performed TAF analyses for telomeric DNA damage but did not find a significant difference in cells expressing this feature of senescence within the callus region with AP treatment (Supplementary Figure 2E, F). These findings thus indicate that targeted clearance of p16+ cells does not significantly improve initial fracture healing but may have a marginal effect at a later stage (e.g., day 28, reflected by a higher AUC under the fracture healing curve, Supplementary Figure 2C) when p16+ cells are more prevalent (Figure 1E).

### p21+ cell clearance suppresses OCH- and neutrophil-derived factors driving osteoclast recruitment and inhibition of bone formation

To investigate how clearance of p21+ cells enhanced fracture healing, we performed single-cell CyTOF phenotyping of callus cells from *p21-ATTAC* mice following clearance of p21+ cells using AP (Figure 3A). Using CITRUS analysis, which generates separately stratified clusters from the original dataset to observe statistical differences (23), we found that AP administration led to downregulation of markers for the SASP, DNA damage, and anti-apoptosis proteins specifically within OCH and neutrophil clusters (Figure 3B-E; Supplementary Figure 3A; note that all changes in Figures 3D, E are statistically significant with FDR < 0.05). Within the OCH cells, CXCL1, which is a potent chemoattractant for neutrophils (24) and osteoclasts (25), as well as TGFβ1, an inhibitor of mineralization and bone formation (11, 26), were significantly reduced by AP treatment (Figure 3D). Other signals suppressed in both OCH cells and neutrophils with AP treatment included IL-1α, known to stimulate osteoclastogenesis (27) and osteoblast apoptosis (28), and STAT1 signaling (measured by pSTAT1 expression), which inhibits fracture-related bone formation (29).

**Figure 3:**
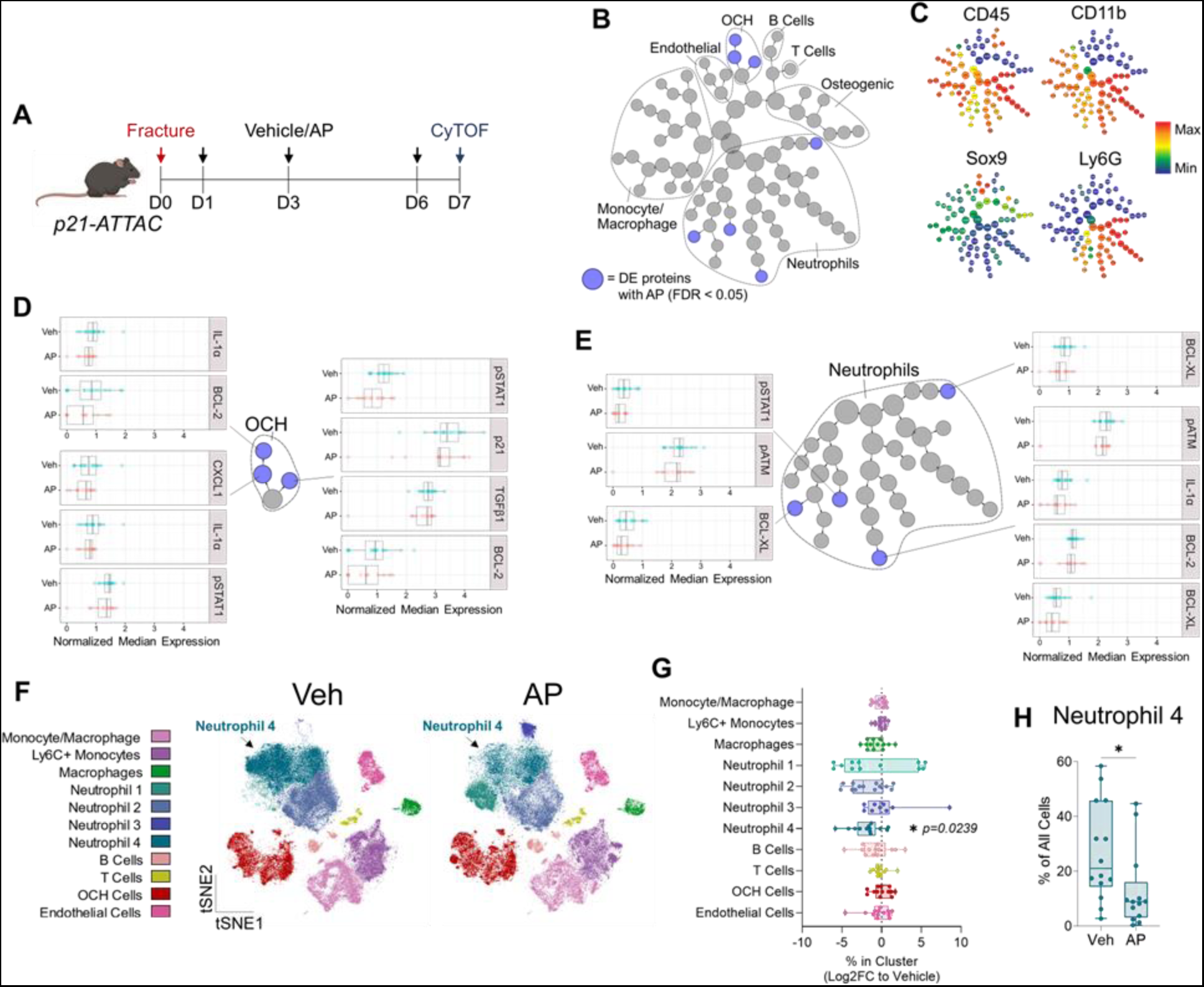
p21+ cell clearance suppresses factors driving osteoclast recruitment and inhibition of bone formation through targeting OCH cells and neutrophils. (A) Schematic outlining CyTOF analysis of callus cells after p21+ cell clearance in *p21-ATTAC* mice. (B) CITRUS analysis reveals reduced expression (blue dots; FDR <0.05) of proteins in OCH and Neutrophil cell clusters. (C) CITRUS expression plots for key identification markers. (D) CITRUS results of differential expression between vehicle- and AP-treated groups in OCH (D) and Neutrophil (E) clusters; all FDR < 0.05. (F) t-SNE visualization and FlowSOM clustering of callus cells from p21-ATTAC mice treated with either Vehicle (Veh) or AP. (G) Quantification of changes in cell cluster percentages after AP treatment (Log2 Fold-change; two-way ANOVA with Sidak multiple comparison test). (H) Quantification of absolute changes in the Neutrophil-4 cluster after AP treatment (Mann-Whitney test). n=16 Vehicle-treated mice (8 female, 8 male), n=13 AP-treated mice (7 female, 6 male).

FlowSOM clustering of these cells (Supplementary Figure 3B, C) demonstrated a significant reduction in a subset of neutrophils (Neutrophil-4) of over 50% after AP treatment (Figure 3F-H) (vehicle: 27% of total cells vs AP: 13%; p=0.0125), but no alterations in the abundance of OCHs or any other cell populations (*e.g.,* macrophages, B- or T-cells). In summary, we found that clearance of p21+ cells in the *p21-ATTAC* mice reduced the numbers of a specific sub-set of neutrophils, did not reduce numbers of inflammatory OCH cells, but did reduce the SASP of the inflammatory OCH cells, including factors related to osteoclast recruitment and suppression of bone formation.

### scRNA-seq analysis of inflammatory p21+ callus cells

Given the critical role of p21+ cells in modulating fracture repair, we next used scRNA-seq to further investigate the inflammatory profile of these cells. To enrich for p21+ cells, we used a reporter mouse in which a validated fragment of the p21 promoter (17) was placed upstream of GFP (Figure 4A). This allowed for FACS isolation of p21+ (GFP+) and p21- (GFP-) callus cells at 14 days post-fracture (Figure 4A). We first validated the increase in GFP+ cells within the fracture site using qRT-PCR and found a significant enrichment of the GFP signal within the callus (Figure 4B). We also performed further qRT-PCR analysis of the GFP+ vs. GFP-cells and detected an increase of SASP markers (*Cxcl2*, *Vegfa* and *Tnfa*) in the GFP+ population (Figure 4C). We then performed scRNA-seq followed by unbiased clustering, leading to 16 distinct clusters (Figure 4D). A number of SASP markers were enriched in p21+ cells, including *Cxcl2, Ccl4, Il1b*, as well as *Tgfb1* (Figure 4E), which has recently been shown to impair fracture healing (11). Consistent with our CyTOF data, the majority of p21+ cells were found within OCH cells (Figure 4F). Due to an increase in clustering variables compared to CyTOF, the scRNA-seq clustering further subgrouped the OCH cells into 2 populations (OCH1 and OCH2), with the OCH1 cells being perhaps earlier in the differentiation lineage (Supplementary Figure 4A, B). Notably, the OCH1 cells also had the highest SASP profile as demonstrated by SenMayo (30) gene set enrichment (Figure 4G). Moreover, the OCH1 cells were predicted to have the highest outgoing strength of all signaling pathways among callus cell types (Figure 4H, I), consistent with a highly secretory profile, particularly in TGFβ signaling (Figure 4J). In addition, OCH1 cells highly expressed activin A (encoded by *Inhba*), a TGFβ superfamily member recently found to mark a distinct proliferative progenitor cell population in the fracture callus (Supplementary Figure 4C) (31).

**Figure 4:**
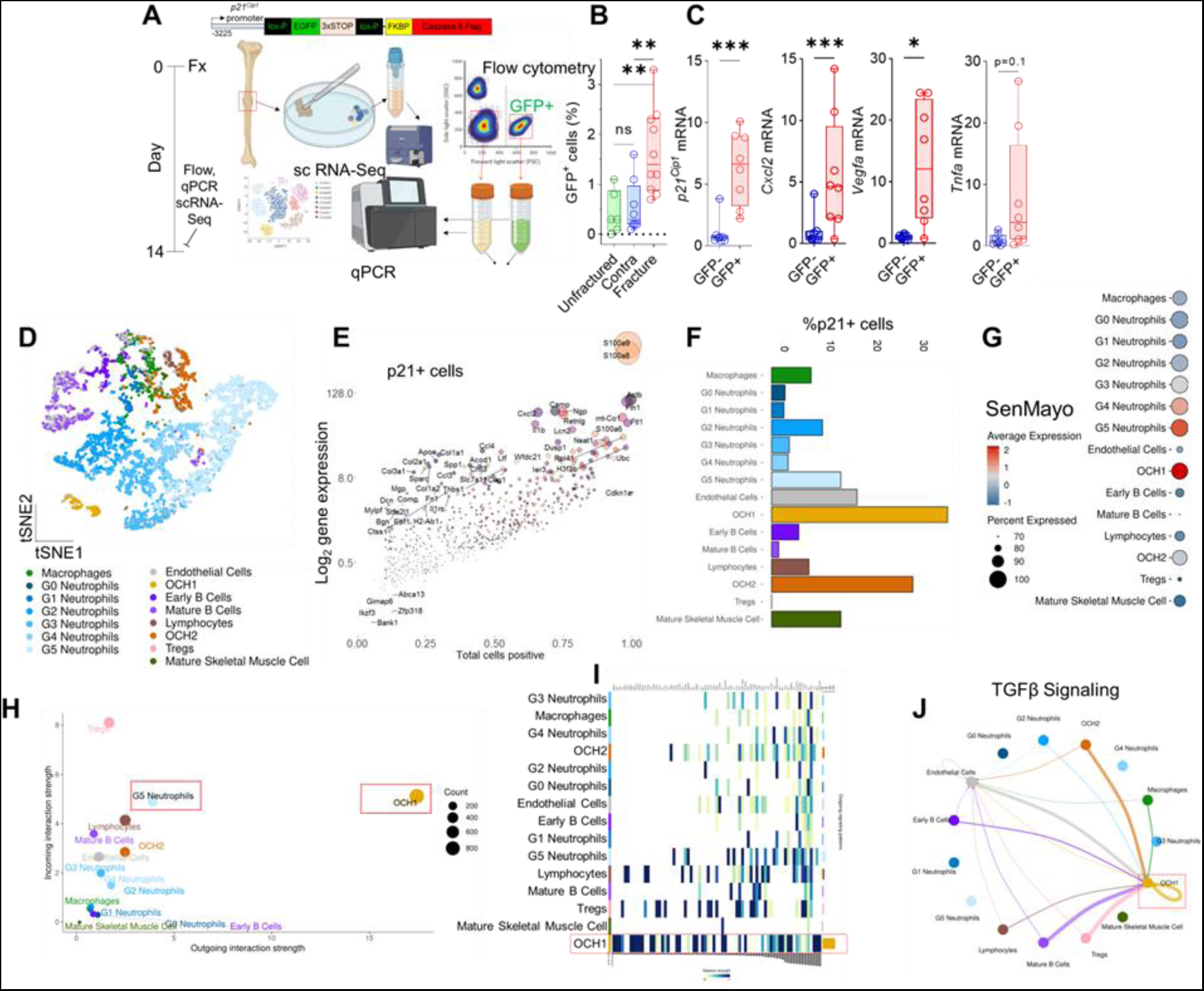
p21+ callus cells are largely highly secretory OCH cells and mature neutrophils. (A) Schematic of p21+ cell isolation and scRNA-sequencing using the p21-reporter mice. (B) GFP^+^ cells were significantly higher in the fractured compared the unfractured contralateral side and an unfractured mouse tibia. (C) *p21^Cip1^*, *Cxcl2*, *Vegfa* and *Tnfa* mRNA expression was significantly enriched in GFP+ cells. (D) scRNA-Seq analysis was performed on 5,994 total callus cells from n=4 mice (n=2 male, n=2 female). (E) Differentially upregulated mRNA transcripts in p21+ cells. (F) Proportion of p21+ cells and (G) SenMayo gene enrichment analysis among clustered cell populations. (H, I) Predicted secretory strength relationships in callus cells among all signaling pathways and (J) TGFβ signaling by CellChat. B: One-Way ANOVA with Tukey’s correction: n=10 fractured (n=4 female, n=6 male), n=8 contralateral sides (n=4 female, n=4 male), n=6 unfractured (n=3 female, n=3 male). (C) GFP^+^ vs. GFP^−^: Mann-Whitney test n=8 mice (n=4 male, n=4 female). *p<0.05, **p<0.01, ***p<0.001.

We also clustered the neutrophils into G1-G5 based on a previously published mouse neutrophil atlas (Figure 4D) (32). The G5 neutrophils had the highest percentage of p21+ cells among the 5 neutrophil populations (Figure 4F), had the second-highest outgoing signal interaction strength among all cell types (Figure 4H), were Ki67- (Supplementary Figure 4D), and were the neutrophil population with the highest SenMayo and ROS score (Supplementary Figure 4E, F). G5 neutrophils are a mature neutrophil population predicted to arise from the peripheral blood (32). Note that due to the greater resolution of the scRNA-seq analysis as compared to CyTOF, we cannot unequivocally equate the G5 population with one of the 4 neutrophil populations identified above by CyTOF. However, like G5 neutrophils, the Neutrophil-4 population that was cleared with AP treatment contained the largest percentage of p21+ cells among all neutrophil subpopulations (Supplementary Figures 4G, H). This suggests that the Neutrophil-4 population identified by CyTOF contains, or is highly enriched for, the G5 neutrophils found by scRNA-seq. This inflammatory neutrophil population was predicted to utilize the thrombospondin1 (THBS1)-Cd47 pathway for communication with the OCH1 population (Supplementary Figure 4I, J). This suggests secretion of THBS1 by the G5 neutrophils, which binds to the CD47 receptor on the OCH1 population, an interaction known to induce paracrine senescence in mesenchymal cells (33, 34).

Next, we aimed to identify regulatory units for the OCH cells. Using SCENIC (35), we reconstructed a regulon-network and, based on AUC-covering on gene density regions, we identified Kruppel-like factor 4 (*Klf4)* to be the most prominent transcription factor for the OCH cells (Supplementary Figure 5A, B). Importantly, *Klf4* has been shown to increase in response to inflammatory stimuli and mediate pro-inflammatory signaling and was of particular importance in the OCH1 and OCH2 clusters (Supplementary Figure 5C) (36).

### Evidence for the appearance of similar injury-related senescent-like cells following muscle injury

To evaluate whether similar, inflammatory p21+ mesenchymal cell populations may be present not only following skeletal fracture, but also following injury across different tissues, we analyzed publically available scRNA-seq data following muscle injury (12). A mesenchymal progenitor cell population in muscle highly analogous to the OCH cells in bone are fibro-adipogenic progenitors (FAPs) (37). Interestingly, following muscle injury, FAPs not only expressed p21 (Supplementary Figure 6A, B), they also demonstrated the highest expression of SASP-associated genes (Supplementary Figure 6C). Similar to the OCH1 cells, FAPS had the highest predicted outgoing signaling strength (Supplementary Figure 6D, E), accompanied by strong TGFβ-signaling to surrounding cells (Supplementary Figure 6F), including activin A expression (Supplementary Figure 5G). Of note, the neutrophil population, which was relatively small in this dataset and thus could not be further subdivided, nonetheless also showed a communication pattern similar to the G5 population towards the OCH1 cells, consisting of the crucial THBS1-CD47 axis from neutrophils to FAPs (Supplementary Figure 6H, I).

### p21+ OCHs are non-growth-arrested inflammatory callus cells

OCHs exhibited the predominant senescence signature of all callus cells, including the highest percentage of p21+ cells and the highest expression of the SenMayo panel, yet it remained unclear why these cells were not reduced after targeted clearance in the *p21-ATTAC* mice. Surprisingly, although OCH1 cells demonstrated a clear senescent-like phenotype by scRNA-seq, they were found to have a high enrichment for proliferation-associated gene expression (Figure 5A). Moreover, out of the two p21+ cell populations shown to be affected by p21+ cell clearance by CyTOF, OCH cells were substantially higher in % Ki67+ cells and Ki67 mean expression as compared to the Neutrophil-4 cells (Figure 5B). Paradoxically, OCH cells still demonstrated an otherwise clear senescent phenotype, demonstrating higher expression of senescence (p21, p53), DNA damage (pATM), and detrimental SASP (IL-1β, PAI-1, TGFβ1, pSTAT1) markers than Neutrophil-4 cells (Figure 5C).

**Figure 5:**
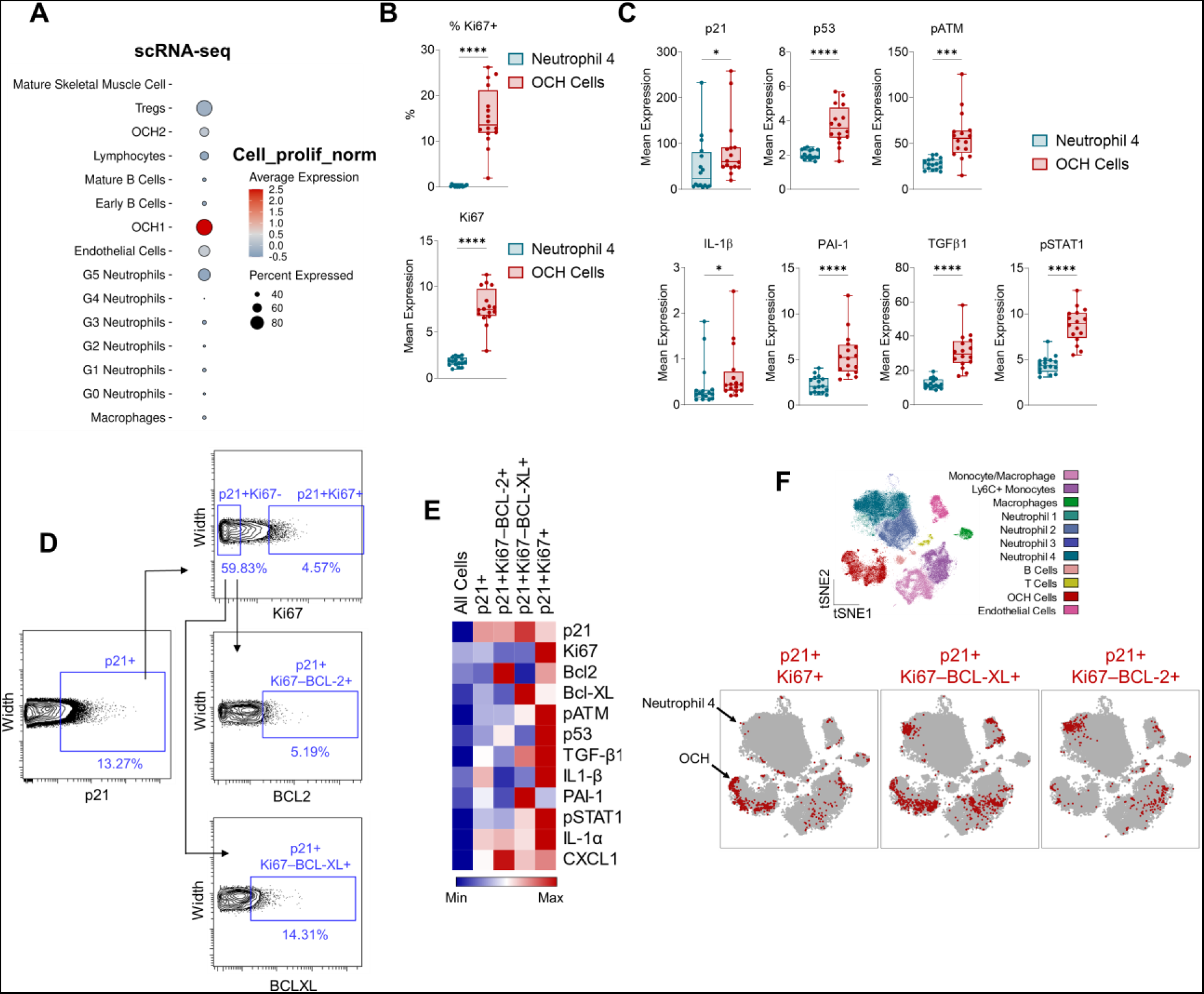
OCH cells are a proliferative population with a senescent-like phenotype. (A) Expression of cell proliferation geneset (GO_0008283) among callus cell populations identified by scRNA-seq (see Figure 4). (B) % Ki67+ and Ki67 mean expression between OCH cell and Neutrophil-4 callus cell clusters identified by CyTOF (see Figure 3). (C) Senescence-related and SASP protein expression between OCH cells and Neutrophil-4 clusters. (D) Gating strategy for p21+/Ki67+, p21+Ki67-BCL2+, and p21+Ki67-BCL-XL+ cell populations in fractured mice. (E) Heatmap demonstrating mean expression of senescence-associated proteins in p21+ subsets by CyTOF. (F) t-SNE visualization of FlowSOM-clustered callus cells overlaid with p21+/Ki67+, p21+Ki67-BCL2+, and p21+Ki67-BCL-X Lcells (red). (A) n=4 mice, (B-F) n=16 mice. (B, C) Mann-Whitney or unpaired t-test, as appropriate. *p<0.05, **p<0.01, ***p<0.001, ****p<0.0001.

We have previously found that, in the context of aging, skeletal cells marked by the senescence-associated cell cycle proteins p21 and p16 contain both senescent and non-senescent (i.e., non-growth-arrested) cell types, with senescent subsets defined as Ki67- and positive for apoptosis-resistance proteins (14). In our callus cells, we subdivided p21+ cells by proliferative (Ki67+) *versus* non-proliferative (Ki67-) cells, and then further divided Ki67-cells into subsets positive for senescence-associated apoptosis-resistance proteins BCL-XL or BCL2 (Figure 5D). While we found that each population demonstrated an enriched SASP, the p21+Ki67+ subpopulation surprisingly exhibited the highest levels of a majority of senescence-associated markers in our CyTOF panel (Figure 5E). These p21+Ki67+ cells were highly enriched in OCH cells (Figure 5F, left panel). By contrast, the p21+Ki67-BCL-2+ cells appeared predominantly within Neutrophil-4 cells (Figure 5F, right panel), while p21+Ki67-BCL-XL+ cells appeared in both Neutrophil-4 and OCH cell clusters (Figure 5F, middle panels). Thus, the predominant inflammatory mesenchymal p21+ cell population in fracture healing is a previously uncharacterized p21+Ki67+ OCH population, rather than truly senescent, growth-arrested cells. However, the appearance of a neutrophil sub-population expressing features of senescence (upregulation of p21, growth arrest, upregulation of BCL2 and/or BCL-XL) was unexpected, although perhaps not unprecedented, as a similar neutrophil population expressing features of senescence has recently been described in the setting of prostate cancer (38).

### The p21+ inflammatory subpopulation of OCHs demonstrates a skeletal stem cell expression profile and is injury-specific

To investigate the OCH population further, we performed additional CyTOF analyses using an expanded antibody panel to include markers known to label skeletal stem cells (SSCs) (CD51 (39), Ctsk (40), LeptinR (41)), chondrogenic cells (Col2a1 (42), NFATc1 (43)), and non-stem stromal cells (Thy1 (44), Embigin (45)). In isolated callus cells, we found 3 separate OCH clusters (positive for Sox9, or Sox6) that expressed markers belonging to distinct stages of differentiation: OCH-Stem (CD51+, LeptinR+, PDGFRα+, Ctsk+, Thy1-), OCH-Mid (CD51+, Thy1+), and OCH-Mature (CD29+, CD200+, Embigin+, ALPL+) (Figure 6A, B). Interestingly, the majority of senescence markers – including those found to be expressed in OCH cells in the previous analyses described above – were most enriched in OCH-Stem cells (Figure 6C, D). Moreover, these cells were overall highest in Ki67 expression, and among the highest in pATM expression, suggesting a DNA-damaged, yet proliferative phenotype (Figure 6E). This OCH-Stem population appeared to be injury-specific, as various manually-gated Sox9+ subsets expressing OCH-Stem SSC markers were highly upregulated in the fracture callus, yet existed at extremely low or non-existent levels in unfractured control mice (Figure 6F, G). This injury-specific OCH-stem population also demonstrated clear upregulation of pATM, TGFβ1, and BCL-XL compared to non-injured controls (Figure 6H). Overall, these data define an injury-related senescent-like OCH population expressing SSC markers that are expanded upon bone fracture and develop features of cellular senescence solely within the context of skeletal injury.

**Figure 6:**
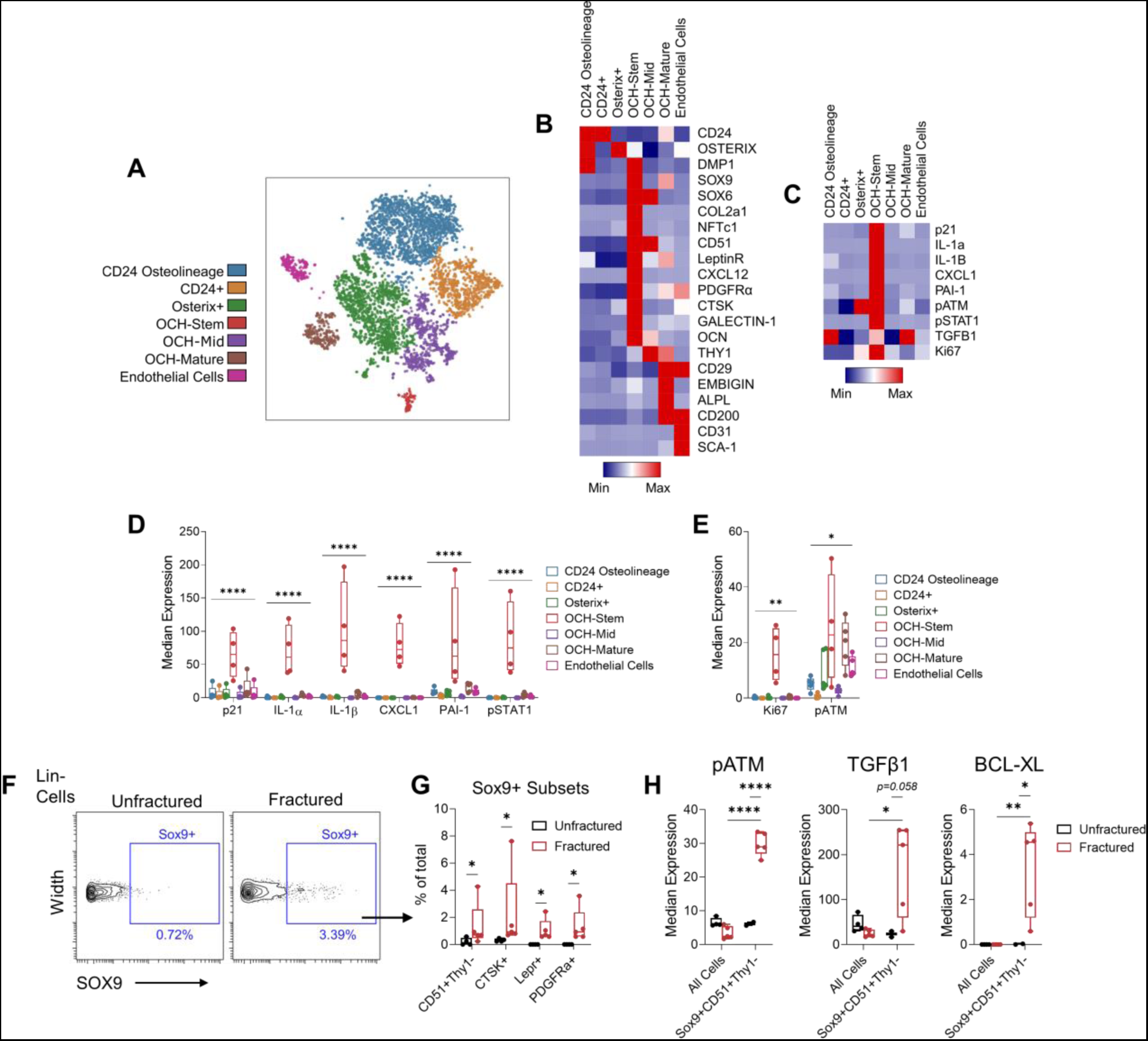
OCHs expressing SSC markers define an injury-specific senescent-like population. (A) t-SNE visualization of FlowSOM clustered callus cell populations collected from fractured wild type C57BL/6 mice. (B, C) Heatmap representation of mean protein expression of (B) identity and (C) senescence-associated markers. (D, E) Quantification of senescence-associated proteins among all clustered cell populations. (F) Gating strategy for Lin-Sox9+ cells in CyTOF samples isolated from either unfractured or fractured bones from young (4-6 month-old) mice. (G) Quantification of manually-gated OCH-Stem clusters in both unfractured and fractured samples. (H) Quantification of senescence-associated proteins in manually-gated OCH-Stem (Lin-Sox9+CD51+) cell populations compared to all cells in both unfractured and fractured samples. n=5 fractured, n=4 unfractured, all female. (D, E, H) Two-way ANOVA with Tukey’s multiple comparisons test, (G) Mann-Whitney test. *p<0.05, **p<0.01, ***p<0.001, ****p<0.0001.

### p21+ cell clearance has no greater effect on fracture healing in aged mice and does not alleviate age-related bone loss

As senescence is clearly linked to aging, we next evaluated if these inflammatory p21+ cells appear in the inflammatory setting of aging and contribute to age-related bone loss. *p21-ATTAC* mice were thus aged to 20-months and treated with either vehicle or AP for 4 months to assess skeletal phenotypes at 24-months of age (Figure 7A), a treatment regimen previously shown to alleviate age-related bone loss in *p16-INK-ATTAC* mice (19). Upon reduction of p21+ cells, indicated by lower *Cdkn1a* mRNA expression in bone (Figure 7B), there were no beneficial changes in any skeletal parameter at any of the three sites examined (femur diaphysis, metaphysis, or lumbar spine) (Figure 7C-H). The few statistically significant changes that were identified were, in fact, detrimental effects on femoral skeletal parameters (Figure 7D, F, G).

**Figure 7:**
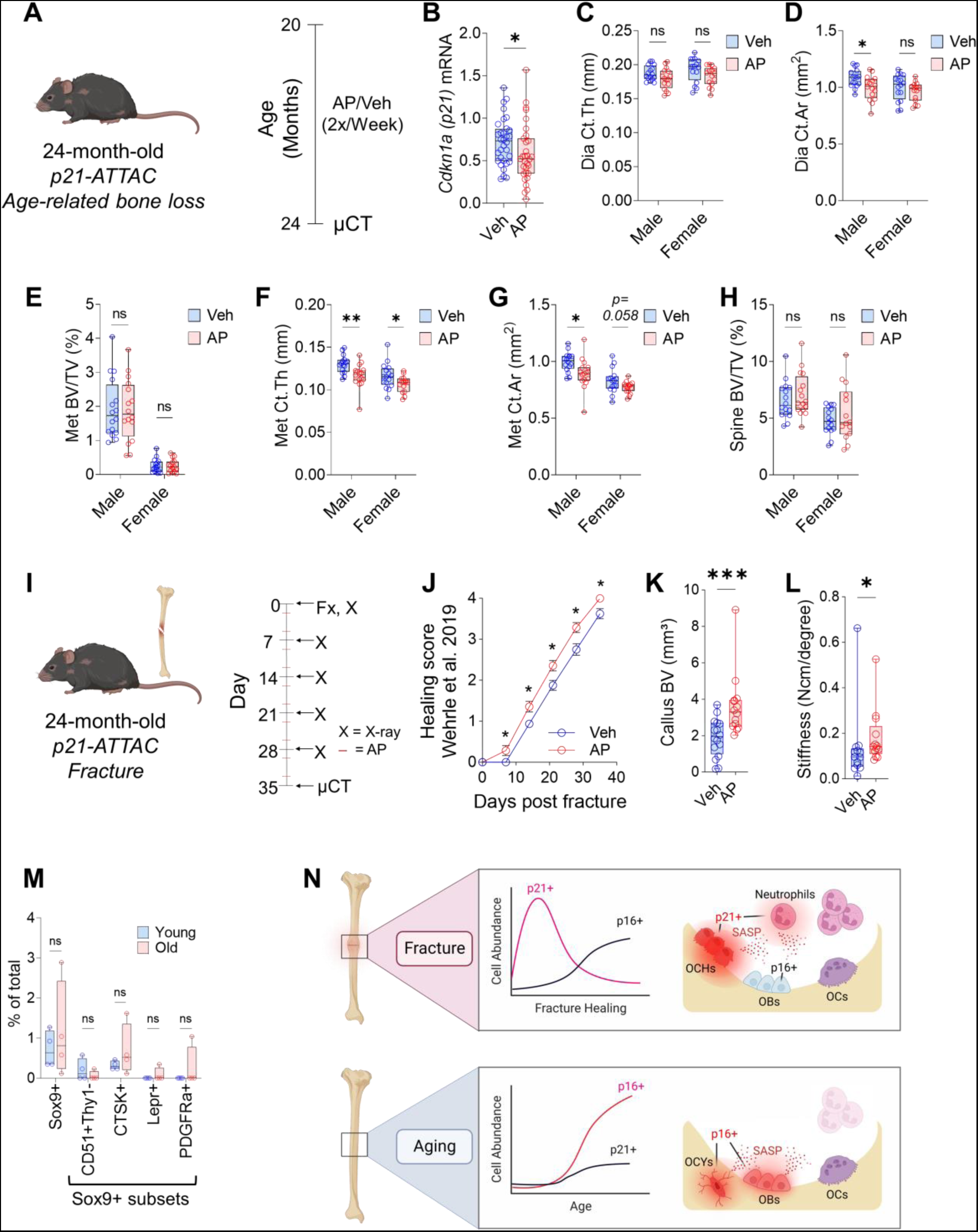
Detrimental effects of p21+ cells on bone metabolism are aging-independent. (A) p21+ cell clearance was performed in old *p21-ATTAC* mice treated at 20 months of age for 4 months with either vehicle (n=32 mice: 16 male, 16 female) or AP (n=32 mice: 16 male, 15 female) twice weekly until sacrifice at 24 months. (B) qPCR measurement of *p21^Cip1^* mRNA expression after AP treatment at sacrifice. (C-G) Skeletal parameters measured at the femur by Micro-CT: (C) diaphyseal (Dia) cortical thickness (Ct.Th), (D) Dia cortical area (Ct.Ar), (E) metaphyseal (Met) trabecular bone volume per total volume (BV/TV), (F) Met Ct.Th, and (G) Met Ct.Ar. (H) Trabecular BV/TV measured at the L5 lumbar vertebra. (I) Schematic for clearance of p21+ cells in old mice undergoing fracture repair. 24-month-old *p21-ATTAC* mice were used to selectively clear p21+ cells through treatment with either vehicle (n=16 mice; 8 male, 8 female) or AP (n=14 mice; 7 male, 7 female) twice weekly over a 5-week fracture healing timecourse. (J) Fracture healing score measured by weekly x-ray. (K) Micro-CT of callus bone volume. (L) Tibial stiffness measured by biomechanical testing (n=13 Veh: 6 male, 7 female. n=13 AP: 7 male, 6 female). (M) CyTOF analysis of OCH-Stem population abundances among CD45-Lin-non-immune cells isolated from the digested hindlimbs of young (6-month-old) and old (24-month-old) wild-type C57BL/6 mice (n=4 mice per group, all female). (N) Schematic of results from injured young bone versus intact aging bone, demonstrating the critical role of transient p21+ cells in delaying fracture healing, but not age-related bone loss; conversely, p16+ cells play a limited role in modulating fracture healing, but alleviate age-related bone loss. OCHs = osteochondroprogenitors, OBs = osteoblasts, OCs = osteoclasts, OCYs = osteocytes. (B-H, K-M) Mann-Whitney or unpaired t test where appropriate, (J) Multiple unpaired t-tests with FDR Correction. *p<0.05, **p<0.01, ***p<0.001.

To test if aging perhaps exacerbates the effect of p21+ cells on impairing fracture healing, we fractured 24-month *p21-ATTAC* mice and treated them with vehicle or AP for 5 weeks postfracture (Figure 7I), identical to our experiments in young mice (Figure 2). Although there was still a beneficial effect of p21+ cell clearance on fracture healing, neither the acceleration (x-ray healing score) nor the endpoint effect (callus bone volume) was any greater in the aged *versus* the young mice (Figure 7J-L). Using CyTOF on bone samples from 24-month-old mice, we found that there was no age-related increase in any of the OCH-Stem subpopulations found to express features of senescence in the setting of fracture (Figure 7M). This suggests that the detrimental effects of senescent-like p21+ cells on bone metabolism are independent of aging. In summary, clearance of p21+ cells in aged mice had no greater effect on fracture healing than in young mice and, in contrast to clearance of p16+ cells (19, 46), did not alleviate age-related bone loss.

## DISCUSSION

In the present study, we identify and define the functional role of cells expressing features of senescence during tissue repair, using fracture healing as our model. These injury-related senescent-like cells, which consist principally of p21-expressing OCH cells and neutrophils, have similarities as well as some key differences from classical senescent cells associated with aging. Specifically, in the context of aging, these cells appear to be principally, if not exclusively, of mesenchymal origin (14, 47). Indeed, cellular senescence was originally defined by Hayflick and Moorhead for mesenchymal cells (48), and whether immune cells express the full features of classical senescent cells remains unclear. However, following fracture, there clearly is a sub-population of neutrophils (Neutrophil-4 by CyTOF, G5 Neutrophils by scRNAseq) that express p21, are growth-arrested, have evidence of DNA damage, express a SASP, and upregulate BCL2 and/or BCL-xL. A second population of these injury-related senescent-like cells consists of mesenchymal OCH cells (OCH-Stem) that also expresses p21, DNA damage markers, and a SASP, but also expresses proliferation markers (p21+Ki67+ cells by CyTOF, p21+ cells positive also for proliferation genes by scRNAseq), in contrast to the growth-arrested classical senescent cells we previously identified in the context of skeletal aging (14). In addition, while classical senescent cells associated with aging are characterized by their persistence over time, the injury-related senescent-like cells are relatively transient, as our previous studies demonstrated that the TAF+ cells in the fracture callus had largely disappeared by day 28 following fracture (10). Finally, although our studies focused on skeletal injury, our demonstration of p21-expressing mesenchymal FAPs following muscle injury with features very analogous to the OCH cells suggests that these injury-related mesenchymal progenitor senescent-like cells appear following not just fracture but following injury across tissues.

Our data further demonstrate that clearance of p21+ cells using a highly specific genetic approach (*p21-ATTAC*) (17) accelerates fracture healing. Moreover, although previous studies demonstrated the beneficial effects on fracture healing of clearing senescent-like cells using a pharmacological approach (D+Q) (10, 11), the present study provides both a clear identification of these senescent-like cells and dissects the relative contributions of the p21 vs. the p16 pathways in driving these injury-related senescent-like cells during fracture healing.

While clearance of p16-expressing senescent cells with aging prevents age-related bone loss (19, 46), it appears that the transiently senescent-like cells following bone injury are largely defined by expression of p21. These findings are entirely consistent with our previous work on focal radiation therapy (17), suggesting that p21 drives injury-specific senescence in the skeleton. Collectively, these findings lead to the hypothesis that age-related bone loss is primarily driven by p16+ cells, whereas acute bone loss following radiation or impaired healing following skeletal injury is principally driven by p21+ cells (Figure 7N). In the case of fracture, the p21+ senescent-like cells are principally OCH cells and a subset of neutrophils while in the context of aging, the p16+ senescent cells consist predominantly of osteocytes (46) and a CD24+ osteolineage cell population (14) (Figure 7N). We should note, however, that findings from these *ATTAC* models need to be further corroborated by complementary approaches not relying on *ATTAC*-mediated clearance of p16+ or p21+ cells. For example, studies using inducible deletion of *p16^Ink4a^* and/or *p21^Cip1^* specifically following fracture or in old mice may, in contrast to *ATTAC*-mediated clearance of senescent cells, prevent the formation of injury- or age-related senescent cells in the first place and help validate findings from the *ATTAC*-clearance models. In addition, further studies are needed to evaluate whether this dichotomy between p16 and p21 as it pertains to aging- vs. injury-induced senescence is specific to the skeleton or is also true for non-skeletal tissues.

As noted above, our CyTOF analyses, complemented by scRNA-seq data, also provide further insights into specific cell populations expressing features of senescence and their interactions. In particular, we found that early (days 1-3) following fracture, there was the appearance of a specific sub-population of neutrophils (G5, based on aligning our scRNA-seq data with a published mouse neutrophil atlas (32), likely contained in the Neutrophil-4 population identified by CyTOF) that is comprised of mature/aged neutrophils that are highly inflammatory (32). Consistent with our findings, previous reports have identified neutrophils as early components of the fracture callus and depletion of neutrophils (*e.g.,* using Ly-6G-antibody treatment) actually impairs fracture healing (49, 50). Our findings demonstrate, however, that a subset of these neutrophils begin to express features of cellular senescence, including increased p21 expression, growth arrest, and a pro-inflammatory SASP, along with expression of genes related to ROS pathways. Moreover, genetic reduction in these p21+ neutrophils, with preservation of non-p21-expressing neutrophils, leads to enhanced fracture repair. Thus, specifically targeting the injury-related senescent-like neutrophils may allow the beneficial effects of non-senescent neutrophils to accelerate repair.

Our demonstration of neutrophils expressing features of senescence is not unprecedented, as Bancaro et al. (38) recently described a very similar population of neutrophils expressing senescent markers and persisting in the tumor microenvironment. In particular, these tumor-infiltrating senescent-like neutrophils also expressed p21 and were highly inflammatory, particularly for SASP factors (38). Moreover, similar to our finding that clearance of senescent-like neutrophils accelerated fracture healing, these investigators also found that both genetic and pharmacological elimination of these tumor-infiltrating senescent-like neutrophils decreased tumor progression in different mouse models of prostate cancer (38). Thus, neutrophils expressing senescent-like features may be part of a broader tissue response to injury and/or cancer. It is also important to note that while terminally differentiated neutrophils are growth-arrested (51), only a sub-set (maximum of ~20% either by CyTOF or scRNAseq in our data) are p21+ but importantly, it is these p21+ neutrophil sub-sets (*i.e.,* Neutrophil-4 by CyTOF or G5 Neutrophils by scRNAseq) that express senescent-like features, including high levels of SASP factors, and are reduced following AP treatment in the *p21-ATTAC* mice.

Our studies also point to important cross-talk between the G5 neutrophils and OCH cells. Previously, the Passos laboratory demonstrated that neutrophils cause telomere dysfunction in neighboring mesenchymal cells in a ROS-dependent manner (52). Moreover, senescent cells mediate the recruitment of neutrophils to their niche, potentially leading to the spread of senescence to surrounding cells (52). Thus, there appears to be a feed-forward loop between G5 neutrophils and OCH cells in the fracture callus whereby the OCH cells recruit inflammatory neutrophils (*e.g.,* through their high expression of CXCL1, a potent chemoattractant factor for neutrophils (24)) that are high in ROS pathways, which causes DNA damage in the OCH cells, leading to a senescent phenotype, which then further recruits neutrophils. Consistent with this, our CellChat analyses found evidence for extensive cross-talk specifically between the G5 neutrophils and OCH cells.

In terms of the OCH cells, we were able to segregate these, using both scRNA-seq and CyTOF, into populations at multiple stages of differentiation. In contrast to the growth-arrested (Ki67-), p21+ neutrophil population that was cleared by AP treatment in the *p21-ATTAC* mice, the OCH cells were predominantly p21+/Ki67+ and their abundance remained unchanged following AP treatment. Rather, these cells had a highly inflammatory phenotype, and their SASP was reduced by AP treatment concomitant with reduction of the p21+ neutrophils. It may be that these cells are initially cleared upon AP treatment yet appear unchanged, as the CyTOF analyses were done 48 hours following the AP dose, due to their rapid re-expansion as a result of their proliferative nature. This may explain why we observed a reduction within the Neutrophil-4 cluster (p21+Ki67-), but not the OCH cells (p21+Ki67+) in mice intermittently cleared of p21+ cells. These p21+Ki67+ cells may represent an inflammatory “pre-senescent” population at the intersection of senescence-associated growth arrest that we (14) and others (53) have recently identified both *in vivo* and *in vitro*. Clearly, the relationship of these cells to fully senescent, growth-arrested cells remains to be further clarified.

The identification of *Klf4* as a key transcription factor within the detrimental OCH cells highlights a therapeutic opportunity. Recently, a small molecule was developed that inhibited the methylation of KLF4 and subsequently downregulated KLF4-mediated gene transcription (54). According to our analyses, this inhibitor might be a promising candidate for accelerating bone healing by impairing the development and/or function of these senescent cells. Moreover, in our scRNA-seq data, OCH cells expressing high levels of p21 and SASP markers were also expressing high levels of *Tgfb1* and based on Cellchat, communicating via the TGFβ signaling pathway with other mesenchymal cell populations. These findings are entirely consistent with the work of Liu et al. (11) who found that senescent callus cells expressed high levels of TGFβ1 and neutralizing antibodies to TGFβ1 prevented the inhibitory effects of these senescent callus cells on the proliferation of MSCs. Collectively, these findings suggest that approaches to inhibit TGFβ1 signaling in the fracture callus may enhance fracture healing, and whether this would be particularly beneficial in the setting of impaired fracture healing (i.e., fracture non-unions) warrants further study.

In order to evaluate whether our findings may extend to tissue injury beyond fracture, we further evaluated p21+ cell populations present following muscle injury using publically available data (12). Remarkably, muscle FAPs, which share multiple characteristics with bone marrow stromal cells and OCH cells (e.g., CD45-CD31-PDGFRα+) (37), were – like the OCH cells –the most highly enriched population for SenMayo genes (30) following muscle injury. FAPs also exhibited very similar cross-talk with neutrophils as we found for OCH cells and neutrophils, in particular through the THBS1-CD47 pathway, which has previously been associated with the induction of paracrine senescence (33, 34). Thus, although further studies across tissues are clearly needed, our findings of a senescent-like, inflammatory mesenchymal progenitor population interacting with infiltrating neutrophils and modulating tissue repair may well extend beyond skeletal injury.

We recognize potential limitations of our work. As noted earlier, we acknowledge that findings from our *ATACC* models need to be corroborated by alternate approaches (*e.g.,* inducible deletion of *p16^Ink4a^* and/or *p21^Cip1^* following fracture or with aging). In addition, while our studies provide interventional evidence, using a genetic clearance strategy, for a causal role for these injury-related senescent-like cells in impairing fracture healing, the specific factors secreted by these cells that modulate fracture repair requires additional studies. As such, while our analyses provide candidate genes and pathways (*e.g.,* the THBS1-CD47 axis from G5 neutrophils to OCH1 cells), fully addressing this issue would require extensive *in vitro* studies involving co-culture of the relevant cell populations along with studies abrogating these pathways *in vivo*. Given the scope of the present work, however, we would submit that these analyses are more appropriate for future work.

In summary, our work combines a comprehensive proteomic, transcriptomic, and highly specific genetic clearance approach to characterize in detail the transiently senescent-like cells that appear following tissue injury. In the case of fracture, we identify these cells as OCH cells and a specific sub-set of neutrophils that express high levels of p21 and a SASP, and we also define important signaling pathways between these key cell populations. Moreover, by directly comparing the effects of genetic senescent cell clearance in *p21-ATTAC vs. p16-INK-ATTAC* mice, we provide further evidence that the injury-related senescent phenotype in the setting of skeletal injury (fracture, radiation (17)) is predominantly driven by p21, particularly in the early healing phase, in contrast to age-related bone loss, which appears to be principally driven by p16 (14, 19, 46). Finally, targeting the injury-related senescent-like populations and/or the pathways we identified may prove beneficial in accelerating fracture healing and potentially repair across multiple tissues (*e.g.,* muscle). In particular, if non-union, which occurs in up to 10% of fractures (55), is shown to be associated with excess accumulation of these senescent-like cell populations, then strategies to reduce or eliminate these cells may provide a novel approach to addressing this vexing clinical problem.

## Materials and methods

### Consideration of sex as a biological variable

Per NIH guidelines (56), we studied both female and male mice. In order to test for possible effects of sex on our primary endpoints, we performed 2-way ANOVA tests on several important parameters of fracture healing (Supplementary Table 3) as recently recommended (57). We found that neither sex alone nor interaction between sex and time was significant, indicating that these cellular effects are not dependent on sex. Thus, both males and females were analyzed together.

### Animal studies

All animal studies were performed under protocols approved by the Institutional Animal Care and Use Committee (IACUC), and experiments were performed in accordance with Mayo Clinic IACUC guidelines. All assessments were performed in a blinded fashion. Mice were housed in ventilated cages and maintained within a pathogen-free, accredited facility under a twelve-hour light/dark cycle with constant temperature (23°C) and access to food (diet details are specified below) and water ad libitum. We used young adult C57BL/6 WT, p16-INK-ATTAC (18), p21-ATTAC (17) mice and transgenic reporter mice with the p21 promoter driving GFP for our experimental procedures. Isoflurane (vaporizer, 1.5-2% in oxygen, inhalation) was used for anesthesia during surgery for induction and maintenance until the surgery was complete (about 15-20 min). Vehicle (Veh) and AP were administered intraperitoneally twice weekly. Vehicle was 4%ETOH 10%PEG-400 and 2%Tween, while AP was dissolved in 4%ETOH, 10%PEG-400 and 2%Tween. The dose was calculated individually with 10mg/kg body weight (BW).

Fractures for all experiments were performed as follows: Mice of comparable mean body weights received a standardized, closed diaphyseal tibial fracture. After a lateral lower leg incision, the left tibia was exposed while the tendons and muscles were protected. A transverse osteotomy with a rotary bone saw was introduced. An Insect Pin (Fine Science Tools, 26001-30, Austerlitz Insect Pin rod diameter 0.03 mm) was inserted retrogradely from the fracture to stabilize the transverse tibial shaft fracture and the distal pin end subsequently directed onto the distal fracture end. After wound closure, postoperative pain management was performed with subcutaneous BupER (0.1 mg/kg body weight [BW]) and the correct position of the pin was immediately affirmed by X-ray. Normal weight bearing postoperatively was allowed.

For the time course study, n=23 male and n=24 female *C57BL/6N WT* untreated mice were used (n=47 in total). For the *p16-INK-ATTAC* study, 6-month-old *p16-INK-ATTAC* mice were used as described in detail elsewhere (19) (Supplementary Figure 2A). Altogether, 73 mice were used: 38 mice for Veh treatment (22 females, 16 males) and 35 for AP treatment (20 female and 15 male mice). For the *p21-ATTAC* study, a total of 47 4-month-old mice were randomized to either vehicle (Veh, n=23; 10 male and 13 female) or AP (AP n=24; 11 male and 13 female) treatment twice weekly (Figure 2A). In addition, fluorescent dyes were subcutaneously injected intraperitoneally to trace newly formed bone: Xylenol orange (0.04mL/animal, 20mg/mL) on day 6, Calcein green (0.1 mL/animal, 2.5mg/mL) at day 13, Alizarin red (0.1mL/animal, 7.5mg/mL) on day 20 and Tetracycline (0.125mL/animal, 5mg/mL) at day 27 post fracture.

### Mouse tissue collection and assessments

Prior to sacrifice, body mass (g) was recorded. The left tibia was stored in 0.9% saline (NaCl) soaked gauze at −20°C for direct *ex vivo* micro-computed tomography (micro-CT) scanning (see Skeletal imaging) and subsequent biomechanical strength testing by standardized torsional testing (see torsional testing). In the *p21-ATTAC and the p16-INK-ATTAC* study, the fractured bone was embedded in methyl-methacrylate and sectioned for fluorescent in situ hybridization (FISH) (see Telomere-Associated Foci [TAF]). In the *p21-ATTAC study*, additional histomorphometrical analyses were performed. In the CyTOF study, the fresh callus was freshly harvested at the indicated timepoints (see CyTOF).

### Cytometry by time of flight (CyTOF) sample preparation

Mice were sacrificed and the left tibia was isolated. Subsequently, the visually verified callus area was removed after cleaning the bone from surrounding tissue. The callus area was minced with a scalpel in FACS buffer. Pieces were then digested three times for 30 minutes each in 0.7mg/ml Collagenase solution (Sigma-Aldrich, Saint Louis, MO]) at 37°C with agitation. In between steps, the solution was filtered over a wet 70µm cell filter, washed three times with PBS and the collected reaction stopped with FBS, while the remaining pieces were further digested. The samples were pooled together, resuspended in RBC lysis buffer for 5 minutes and diluted in FACS buffer. The remaining solution was resuspended and kept on ice.

### CyTOF Antibodies

Metal-conjugated antibodies used in this study are summarized in the Supplementary Table 1. Except commercially available pre-conjugated antibodies (Fluidigm Sciences), all antibodies were conjugated to isotopically enriched lanthanide metals using the MaxPAR antibody conjugation kit (Fluidigm Sciences), according to the manufacturer’s recommended protocol. Labeled antibodies were stored at 4°C in PBS supplemented with glycerol, 0.05% BSA and 0.05% sodium azide. All antibodies were tested with control beads as well as positive and negative control cells. A detailed validation of the key antibodies used (e.g., p21, p16, others) is included in a recent publication from our group (14).

### CyTOF antibody staining and sample processing

Custom conjugated antibodies were generated in-house through the Mayo Clinic Hybridoma Core using Maxpar X8 Ab labeling kits (Standard BioTools) according to the manufacturer’s protocol. Conjugated antibody concentrations were measured by absorbance at OD280nm and then normalized to a 5μg/μL stock concentration. Isolated cells were resuspended in 1 mL of Cell Staining Buffer (CSB) (Standard BioTools) and incubated for 5 minutes with 0.5 µm Cisplatin solution (Standard BioTools) in PBS. Samples were then washed twice with CSB. An antibody cocktail of the entire phenotyping panel (Supplementary Table 1) was prepared as a master mix prior to adding 50 µL of cocktail to samples resuspended in 50 µL of CSB. Samples were then incubated at room temperature for 45 minutes. Samples were washed twice then fixed with 2% PFA (Standard BioTools) in PBS. After fixation and wash, samples were resuspended in 30 nM intercalation solution (Standard BioTools) and incubated overnight at 4°C. On the following morning, cells were washed with PBS and resuspended in a 1:10 solution of calibration beads and cell acquisition solution (CAS) (Standard BioTools) at a concentration of 0.5×10^6^ cells/mL. Prior to data acquisition, samples were filtered through a 35 µm blue cap tube (Falcon). The sample was loaded onto a Helios CyTOF system (Standard BioTools) and acquired at a rate of 200-400 events per second. Data were collected as .FCS files using the Cytof software (Version 6.7.1014). After acquisition, intra-file signal drift was normalized to acquired calibration bead signal using Cytof software.

### CyTOF data analysis

Initial processing and clustering. Cleanup of cell debris–including removal of beads, dead cells, and doublets–was performed (Supplementary Fig. 1A) using Cytobank software(58, 59). Visual representation of single-cell data was achieved using viSNE mapping (5,000 iterations, 100 perplexity, 0.5 theta), which is based on the t-Distributed Stochastic Neighbor Embedding (t-SNE) algorithm.(60) FlowSOM clustering was performed within Cytobank (hierarchical consensus, 10 iterations) and cluster labels were assigned using established literature on skeletal cell types, with relative marker intensities per cluster visualized by heatmap. FCS files were exported, concatenated in R, then re-uploaded for visualization of merged populations. Quantified values were exported to Graphpad Prism 8 to construct plots and perform statistical analyses. CITRUS analyses (61) were performed in Cytobank using Significance Analysis of Microarrays (SAM) correlative association model. Nearest Shrunken Centroid (PAMR) and L1-Penalized Regression (LASSO via GLMNET) predictive association models were run simultaneously to analyze model error rates to confirm validity of the statistical model. For CITRUS assessment of median expression changes, cells were clustered by identification markers and statistics channels included all functional markers; for assessment of abundances, all markers were used for clustering. All CITRUS analyses used the following settings: 2,000 events samples per file, 2% minimum cluster size, 5 cross validation folds, and 5% false discovery rate (FDR).

### Quantitative real-time polymerase chain reaction (qRT-PCR) analysis

For callus analyses, callus and contralateral intact bone were removed as described previously (10) and immediately homogenized in QIAzol Lysis Reagent (QIAGEN, Valencia, CA), and stored at −80°C. In short, the soft tissue was removed, a 7 mm section around the fracture site (and a 7 mm section at the same location on the intact contralateral site) was extracted and homogenized in QIAzol. Subsequent RNA extraction, cDNA synthesis, and targeted gene expression measurements of mRNA levels by qRT-PCR were performed as described previously (62). Total RNA was extracted according to the manufacturer’s instructions using QIAzol Lysis Reagent. Purification with RNeasy Mini Columns (QIAGEN, Valencia, CA) was subsequently performed. On-column Rnase-free Dnase solution (QIAGEN, Valencia, CA), was applied to degrade contaminating genomic DNA. RNA quantity was assessed with Nanodrop spectrophotometry (Thermo Fisher Scientific, Wilmington, DE). Standard reverse transcriptase was performed using High-Capacity cDNA Reverse Transcription Kit (Applied Biosystems by Life Technologies, Foster City, CA). Transcript mRNA levels were determined by qRT-PCR on the ABI Prism 7900HT Real Time System (Applied Biosystems, Carlsbad, CA), using SYBR green (Qiagen, Valencia, CA). The mouse primer sequences, designed using Primer Express Software Version 3.0 (Applied Biosystems), for the genes measured by SYBR green are provided in Supplementary Table 2. Input RNA was normalized using two reference genes (*Actb*, *Gapdh*) from which the most stable reference gene was determined by the geNorm algorithm. For each sample, the median cycle threshold (Ct) of each gene (run in triplicate) was normalized to the geometric mean of the median Ct of the most stable reference gene. The delta Ct for each gene was used to calculate the relative mRNA expression changes for each sample. Genes with Ct values >35 were considered not expressed (NE), as done previously (19).

### Skeletal imaging: Radiographical fracture healing assessment

All imaging and analysis was performed in a blinded fashion as described by our group previously (10). In short, radiographs of the fracture site were taken under anesthesia after surgery and on a weekly basis. Therein, mice were in a supine position and both limbs extended. We assessed both the anteroposterior (ap) and lateral (lat) planes. Radiographs were evaluated by two blinded researchers and scored for fracture healing using the score by Wehrle et al. (20) Quantification of the fracture callus was performed with FIJI (NIH, Bethesda, MD, USA), as described elsewhere (63).

### Skeletal imaging: Ex vivo micro-CT imaging

At the study endpoint, callus volume of the fracture site was performed. Scan settings were: 55 kVp, 10.5 μm voxel size, 21.5 diameter, 145mA, 300 ms integration time. For the callus volume measurement, a threshold of 190 and 450 were chosen according to the manufacturer’s protocols (Scanco Medical AG, Basserdorf, Switzerland).

### Torsional testing of tibiae

Tests of torsional load were performed in a blinded fashion. The pin was removed and the tibia embedded in the tibial pleateau as distal tibia. Subsequently, the torsional load was applied at speed of 5°/second for a maximum of 36 seconds. The primary endpoints were maximum rotation angle at failure (Deg) and stiffness (N-cm/degree). The maximum torque was the highest force that the bone could sustain before fracture, and stiffness was calculated from the linear portion of the loading curve (higher values for both are indicative of stronger bone) (64).

### Bone histomorphometry

All histomorphometric analyses were performed in a blinded manner. For dynamic histomorphometry, mice were injected subcutaneously with Xylenol orange (0.04mL/animal, 20mg/mL), calcein (0.1 mL/animal, 2.5mg/mL), Alizarin Red (0.1mL/animal, 7.5mg/mL) and Tetracycline (0.125mL/animal, 5mg/mL), on days 6, 13, 20 and 27 post fracture, respectively. The healed tibia was embedded in MMA, sectioned, and stained with Masson Trichrome to determine osteoblast numbers/bone perimeter (N.Ob/B.Pm,/mm), or stained for tartrate-resistant acid phosphatase (TRAP) activity to assess osteoclast numbers per bone perimeter, N.Oc/B.Pm,/mm). To quantify the different stages of bone healing, sections were left unstained to quantify trabecular mineralizing surfaces (mineral apposition rate, MAR, µ/d; bone formation rate/bone surface, BFR/BS, µm^3^/µm^2^/d), which were quantified in the center of the former proximal and distal fracture end. Osteoblast (N.Ob/B.Pm) and osteoclast (N.Oc/B.Pm) numbers were assessed at the same place to verify trabecular assessments. All histomorphometric measurements and calculations were performed with the Osteomeasure Analysis system (Osteometrics, Atlanta, Georgia).

In the *p21-ATTAC and p16-INK-ATTAC* studies, both hindlimbs were used for RNA analysis. The callus area was minced with a scalpel in FACS buffer, and homogenized in QIAzol Lysis Reagent (QIAGEN, Valencia, CA), and frozen at −80°C for real-time quantitative polymerase chain reaction (qRT-PCR) mRNA gene expression analyses (see Real-time quantitative polymerase chain reaction analysis). At the contralateral side, the diaphyseal area was prepared in the same manner, with a bone fracture of the same size as the contralateral callus.

### Telomere-associated foci (TAF) assay

TAF was performed on murine hindlimbs of non-decalcified methylmethacrylate (MMA)-embedded sections as previously described (13). Briefly, bone sections were de-plasticized and hydrated in EtOH gradient followed by water and PBS. Antigen was retrieved by incubation in Tris-EDTA (pH 9.0) at 95°C for 15 min. After cool-down and subsequent hydration with water and PBS (0.5% Tween-20/0.1% Triton100X), slides were placed in a blocking buffer (1:60 normal goat serum [Vector Laboratories; Cat. #S-1000] in 0.1% BSA/PBS) for 30 min at RT. The primary antibody targeting γ-H2A.X (Cell Signaling, #9718, 1:200) was diluted in blocking buffer and incubated overnight at 4°C in a humid chamber. On the next day, slides were washed with PBS-TT followed by PBS alone, and then incubated for 30 min with secondary goat, anti-rabbit antibody biotinylated (Vector Laboratories, #BA-1000, 1:200) in blocking buffer. Afterwards, the slides were washed with PBS-TT followed by PBS alone, and then incubated for 60 min with the tertiary antibody (Cy5 Streptavidin, Vector Laboratories, #SA-1500, 1:500) in PBS. The slides were washed three times with PBS, followed by FISH for TAF detection. Following 4% paraformaldehyde (PFA) crosslinking for 20 min, sections were washed three times (five minutes each in PBS), and dehydrated in graded ice-cold EtOH (70%, 90%, and 100% for three minutes each). Sections were dried and denatured for 10 min at 80°C in hybridization buffer (0.1M Tris, pH 7.2, 25mM MgCl2, 70% formamide [Sigma-Aldrich, Saint Louis, MO], 5% blocking reagent [Roche] with 1.0 μg/mL of Cy-3-labeled telomere-specific [CCCTAA] peptide nucleic acid [PNA] probe [TelC-Cy3, Panagene Inc., Korea; Cat. #F1002]). The slides were hybridized in a humid chamber for two hours at RT. After that, the sections were washed and mounted with Vectashield DAPI-containing mounting media (Life Technologies) prior to image acquisition and analysis. The number of TAF per cell was quantified in a blinded fashion by examining overlap of signals from the telomere probe with γ-H2A.X. At least 100 cells were counted per bone. The mean number of TAF per cell in bone diaphysis and/ or callus was quantified using FIJI (Image J; NIH, Bethesda, MD, USA), and the percentage (%) of TAF-positive (TAF+) cells was calculated for each mouse based on the following criteria: % of cells with ≥1TAF, % of cells with ≥2 TAF, and % of cells with ≥3 TAF, respectively. A senescent cell was defined with a cut-off ≥3 TAF per cell.(65)

### scRNA-seq sample preparation

For the scRNA-seq study, we used a total of four mice for single cell sequencing (2 male, 2 female) and a total of n=8 (4 male, 4 female) for flow cytometry and performed surgery as described above for preparation of the samples for CyTOF. The three samples per bone were pooled together and resuspended in RBC lysis buffer for 5 minutes and diluted in FACS buffer. The remaining solution was resuspended and kept on ice, and sorted by FACS.

### Flow cytometry-based sorting of GFP^+^ cells

Following the cell isolation from callus, the cells were live sorted by staining with propidium iodide (PO-PRO I, cat. no.: P3581) in staining buffer. After that, the cells were electronically gated on live GFP^+^ cells based on the wavelength 509nm and a low expression of PO-PRO I (indicating live cells) and subjected to sorting through a FACSAria digital cytometer running FACSDiva v 8.0.1 software (BD Biosciences, Franklin Lakes, NJ, USA). The GFP^+^ and GFP^−^ cells as remaining cells were either kept in QIAzol Lysis Reagent (QIAGEN, Valencia, CA), and stored at −80°C or immediately used for single cell analysis.

### scRNA-seq analysis

After FACS, the single cell suspension was loaded onto the 10x chromium device using Version 2 chemistry. The samples were sequenced at GeneWiz where we targeted 50,000 reads per cell on an Illumina HiSeq 4000 device. We performed two lanes of sequencing using these parameters for each sample. scRNA-seq data were aligned and quantified using the 10X Genomics Cell Ranger Software Suite (v6.1.1) against the murine reference genome (mm10). The Seurat package (v4.1)(66, 67) was used to perform integrated analyses of single cells. Genes expressed in <3 cells and cells that expressed <200 genes and >20% mitochondria genes were excluded from downstream analysis in each sample. The dataset was SCTransform-normalized and the top 3000 highly variable genes across cells were selected. The datasets were integrated based on anchors identified between datasets before Principal Component Analysis (PCA) was performed for linear dimensional reduction. A shared nearest neighbor (SNN) Graph was constructed in order to identify clusters on the low-dimensional space (top 30 statistically significant principal components, PCs). An unbiased clustering according to the recommendations of the Seurat package was used, and a resolution of 1.4 led to 38 distinct cellular clusters. These were manually assigned to 18 cell types (Supplementary Figure 4) The heatmaps were generated in Seurat (v4.1.0) using the top10 differentially expressed genes based on the average log2Foldchange, after applying the FindAllMarkers function (min.pct=0.25, logfc.threshold=0.25). For Uniform Manifold Approximation and Projection for Dimension Reduction (UMAP) calculations, the RunUMAP function (dims = 1:40, reduction = “pca”) was utilized, and both DimPlot (seurat4) and plot_cells (monocle3, v.1.2.0) used for plotting.

For pseudotime analyses, trajectory interference was generated via RNA velocity,(68) monocle3 and monocle2. For RNA velocity, the raw sequencing reads from fastq files were arranged into spliced and unspliced matrices by velocyto. The RNA velocity was the inferred with the stochastic model of Scvelo and Kallisto.(69) Filtering out genes with no more than 20 counts in spliced and unspliced matrices reduced the subsequent Seurat object. After dimensional reduction and PCA- as UMAP calculation, velocity was run onto the PCA reduction revealing the subsequent pseudotemporal trajectory based on spliced and unspliced variants.

For monocle, an independent component analysis (ICA) dimensional reduction is performed, followed by a projection of a minimal spanning tree (MST) of the cell’s location in this reduced space. Each cell is assigned a pseudotemporal space. Monocle 2 was used to preprocess, perform UMAP reduction, and reduce the dimensionality using the DDRTree algorithm with a maximum of four dimensions. Subsequently, the cells were ordered and genes plotted along the reduced dimension. Differential gene testing has been performed with the formula “~sm.ns(Pseudotime)”, and the results were restricted by a qvalue<0.1.(70, 71)

For the signaling network, cellChat (1.1.3) has been utilized aggregating a cell-cell communication network from significant signaling genes and interactions (threshold.p=0.05) according to netAnalysis_signalingRole after the centrality scores were calculated in the inferred intercellular communication network (“netP”, min.cells=10). The regulatory units analysis has been performed using SCENIC (1.2.4).(72) The network analysis was conducted with Cytoscape 3.8.2 and the plugin iRegulon.(73)

### Statistics

Graphical data are shown as Means ± standard error of the mean (SEM) unless otherwise specified. The sample sizes were determined based on previously conducted and published experiments (e.g. Farr et al. (19) and Saul et al. (10)) in which statistically significant differences were observed among various bone parameters in response to multiple interventions in our laboratory. The used animal numbers are indicated in the figure legends; all samples presented represent biological replicates. We did not exclude mice, samples, or data points from analyses. Data were examined for normality and distribution using dot plots and histograms; all variables were examined for skewness and kurtosis. If the normality or equal variance assumptions for parametric analysis methods were not met, data were analyzed using non-parametric tests (e.g., Wilcoxon Rank Sum test, Mann-Whitney U test). For parametric tests, depending on the comparison, differences between groups were analyzed by independent samples t-test or one-way ANOVA, where justified as appropriate. When ANOVA determined a statistically significant (p<0.05) effect, pairwise multiple comparisons were performed and the Tukey post-hoc method was applied unless specified otherwise. Statistical analyses were performed using either GraphPad Prism (Version 9.0) or R version 4.0.2. A p-value <0.05 (two-tailed) was considered statistically significant.

### Study approval

Animal studies were performed under protocols approved by the Institutional Animal Care and Use Committee (IACUC), and experiments were performed in accordance with Mayo Clinic IACUC guidelines.

## Acknowledgements

This work was supported by the German Research Foundation (DFG, 413501650) (D.S.), National Institutes of Health (NIH) grants P01 AG062413 (S.K., J.N.F.), R01 AG076515 (S.K., D.G.M.), R01 DK128552 (J.N.F.), R01 AG063707 (D.G.M.), R01 AG068048 (JFP); UH3 CA268103 (JFP) and R01 AG082708 (JFP), The Glenn Foundation For Medical Research (JFP) R01 AG082681 (A.C), and Mildred Scheel postdoc fellowship by the German Cancer Aid (R.L.K.).

## Competing interests

None

## Data availability

The single cell RNA-Seq data are available at the GEO repository (GSE253863). All additional data is available by the corresponding author on request.

## Author Contributions

DS, MLD, and SK conceived and designed research studies. DS, MLD, JLR, MNF, RLK, SJV, and MR conducted experiments and acquired data. DS, MLD, and SK analyzed and interpreted data with input and advice from N.L., A.C., R. P., J.F.P., and J.N.F. DS, MLD, and SK wrote the manuscript with input and edits from all authors.

## Notes

**Conflict-of-interest statement**: The authors have declared that no conflict of interest exists.

### Competing Interest Statement

The authors have declared no competing interest.

