## Supplementary Tables and Figures for "Osteochondroprogenitor cells and neutrophils expressing p21 and senescence markers modulate fracture repair"

### Saul et al. Supplementary Tables and Figures

**SUPPLEMENTARY TABLES**

**Supplementary Table 1.** CyTOF antibodies, conjugated metals and source.

| **Timecourse Panel** | | | |
|---|---|---|---|
| **Marker** | **Metal** | **Vendor** | **Cat #** |
| CD45 | 089Y | Fluidigm | 3089005B |
| Ki67 | 106Cd | Biolegend | 103051 |
| CD11b | 110Cd | Biolegend | 101201 |
| OCN | 111Cd | Santa Cruz | sc-376835 |
| Bcl-2 | 112Cd | Biolegend | 633502 |
| Ly6C | 113Cd | Biolegend | 128039 |
| p53 | 116Cd | Abcam | ab252388 |
| Ctsk/Cathepsin K | 141Pr | Abcam | ab37259 |
| CD24 | 142Nd | Biolegend | 101802 |
| Dmp1 | 143Nd | ThermoFisher | PA5-47621 |
| SDF-1 / CXCL12 | 144Nd | R&D Systems | MAB350-100 |
| CD115 / CSF1R | 146Nd | Biolegend | 135521 |
| Runx2 | 147Sm | Abcam | ab76956 |
| CD140a / PDGFRα | 148Nd | Biolegend | 135902 |
| CD68 | 149Sm | Biolegend | 137002 |
| TNFRSF11A/CD265 | 150Nd | BioLegend | 119803 |
| SP7 | 151Eu | Invitrogen | PA5-40411 |
| CD200 | 152Sm | Biolegend | 123802 |
| PPARy | 153Eu | Invitrogen | PA3-821A |
| ALPL | 154Sm | R&D Systems | af2910 |
| p16 | 155Gd | Abcam | ab232402 |
| CD73 | 156Gd | BioLegend | 127202 |
| F4/80 | 158Gd | Biolegend | 123101 |
| IL-1β | 159Tb | Cell Signaling | 31202 |
| CD206 | 160Gd | Biolegend | 141702 |
| CX3CR1 | 161Dy | Biolegend | 149002 |
| Sox9 | 163Dy | Abcam | ab220812 |
| CD31 | 165Ho | Fluidigm | 3165013B |
| Sclerostin | 166Er | Abcam | ab63097 |
| ACP5 | 167Er | Abcam | ab83050 |
| Sox6 | 168Er | Biorbyt | orb669202 |
| Ly-6A/Sca-1 | 169Tm | Fluidigm | 3169015B |
| CD169 | 170Er | Fluidigm | 3170018B |
| IL-1α | 171Yb | Biolegend | 503202 |
| E11 | 174Yb | BioLegend | 127401 |
| CXCL1 | 175Lu | R&D Systems | MAB453-500 |
| p21 | 176Yb | Santa Cruz | sc-6246 |

| ***p21-ATTAC* Panel** | | | |
|---|---|---|---|
| **Marker** | **Metal** | **Vendor** | **Cat #** |
| CD45 | 089Y | Fluidigm | 3089005B |
| Ki67 | 106Cd | Biolegend | 350523 |
| CD11b | 110Cd | Biolegend | 101249 |

| | | | |
|---|---|---|---|
| OCN | 111Cd | Santa Cruz | sc-376835 |
| Bcl-2 | 112Cd | Biolegend | 633502 |
| Ly6C | 113Cd | Biolegend | 128039 |
| pATM | 114Cd | Invitrogen | 14-0146-82 |
| p53 | 116Cd | Abcam | ab252388 |
| Ctsk/Cathepsin K | 141Pr | Abcam | ab37259 |
| CD24 | 142Nd | Biolegend | 101802 |
| Dmp1 | 143Nd | ThermoFisher | PA5-47621 |
| TGFB1 | 144Nd | Novus Biologicals | NBP2-74495 |
| CD19 | 145Nd | Biolegend | 115502 |
| CD115 / CSF1R | 146Nd | Biolegend | 135521 |
| Runx2 | 147Sm | Abcam | ab76956 |
| CD140a / PDGFRα | 148Nd | Biolegend | 135902 |
| CD68 | 149Sm | Biolegend | 137002 |
| TNFRSF11A/CD265 | 150Nd | BioLegend | 119803 |
| SP7 | 151Eu | Invitrogen | PA5-40411 |
| CD200 | 152Sm | Biolegend | 123802 |
| PPARγ | 153Eu | Invitrogen | PA3-821A |
| ALPL | 154Sm | R&D Systems | af2910 |
| p16 | 155Gd | Abcam | ab232402 |
| F4/80 | 158Gd | Biolegend | 123101 |
| IL-1β | 159Tb | Cell Signaling | 31202 |
| PAI-1 | 160Gd | Abcam | ab125687 |
| CX3CR1 | 161Dy | Biolegend | 149002 |
| Sox9 | 163Dy | Abcam | ab220812 |
| pSTAT1 | 164Dy | Novus Biologicals | AF2894-SP |
| CD31 | 165Ho | Fluidigm | 3165013B |
| Sox6 | 168Er | Biorbyt | orb669202 |
| Ly-6A/Sca-1 | 169Tm | Fluidigm | 3169015B |
| Ly-6G | 170Er | Biolegend | 127637 |
| IL-1α | 171Yb | Biolegend | 503202 |
| CD206 (MMR) | 173Yb | Biolegend | 141702 |
| SDF-1 / CXCL12 | 174Yb | R&D Systems | MAB350-100 |
| CXCL1 | 175Lu | R&D Systems | MAB453-500 |
| p21 | 176Yb | Santa Cruz | sc-6246 |
| CD3e | 196Pt | Biolegend | 100302 |

| **OCH Detailed Panel** | | | |
|---|---|---|---|
| **Marker** | **Metal** | **Vendor** | **Cat #** |
| CD45 | 089Y | Fluidigm | 3089005B |
| Ki67 | 106Cd | Biolegend | 350523 |
| CD11b | 110Cd | Biolegend | 101249 |
| OCN | 111Cd | Santa Cruz | sc-376835 |
| Bcl-2 | 112Cd | Biolegend | 633502 |
| Ly6C | 113Cd | Biolegend | 128039 |
| pATM | 114Cd | Invitrogen | 14-0146-82 |
| CD51 | 115In | Biolegend | 153202 |
| NFATc1 (NFAT2) | 139La | Biolegend | 649602 |
| Ctsk/Cathepsin K | 141Pr | Abcam | ab37259 |

| | | | |
|---|---|---|---|
| CD24 | 142Nd | Biolegend | 101802 |
| Dmp1 | 143Nd | ThermoFisher | PA5-47621 |
| TGFB1 | 144Nd | Novus Biologicals | NBP2-74495 |
| CD19 | 145Nd | Biolegend | 115502 |
| LeptinR | 146Nd | R&D Systems | AF497 |
| Runx2 | 147Sm | Abcam | ab76956 |
| CD140a / PDGFRα | 148Nd | Biolegend | 135902 |
| SP7 | 151Eu | Invitrogen | PA5-40411 |
| CD200 | 152Sm | Biolegend | 123802 |
| PPARy | 153Eu | Invitrogen | PA3-821A |
| ALPL | 154Sm | R&D Systems | af2910 |
| p16 | 155Gd | Abcam | ab232402 |
| Embigin | 156Gd | ThermoFisher | 14-5839-85 |
| CD29 | 158Gd | Biolegend | 102235 |
| IL-1β | 159Tb | Cell Signaling | 31202 |
| PAI-1 | 160Gd | Abcam | ab125687 |
| Sox9 | 163Dy | Abcam | ab220812 |
| pSTAT1 | 164Dy | Novus Biologicals | AF2894-SP |
| CD31 | 165Ho | Fluidigm | 3165013B |
| BCL-XL | 167Er | Abcam | ab32370 |
| Sox6 | 168Er | Biorbyt | orb669202 |
| Ly-6A/Sca-1 | 169Tm | Fluidigm | 3169015B |
| Ly-6G | 170Er | Biolegend | 127637 |
| IL-1α | 171Yb | Biolegend | 503202 |
| Col2a1 | 172Yb | Abcam | ab34712 |
| Galectin-1 | 173Yb | Abcam | ab225540 |
| SDF-1 / CXCL12 | 174Yb | R&D Systems | MAB350-100 |
| CXCL1 | 175Lu | R&D Systems | MAB453-500 |
| p21 | 176Yb | Santa Cruz | sc-6246 |
| CD3e | 196Pt | Biolegend | 100302 |
| PCNA | 198Pt | Biolegend | 307902 |

**Supplementary Table 2.** Mouse qPCR primer sequences

| Category | Gene | Forward Primer sequence (5'-3') | Reverse Primer Sequence (5'-3') |
|---|---|---|---|
| Senescence | *Cdkn2a*$^{Ink4a}$ | GAACTCTTTCGGTCGTACCC | AGTTCGAATCTGCACCGTAGT |
| Senescence | *Cdkn2a*$^{pan}$ | AGCTCTTCTGCTCAACTACGG | GGAGAAGGTAGTGGGGTCCT |
| Senescence | *Cdkn1a*$^{Cip1}$ | GAACATCTCAGGGCCGAAAA | TGCGCTTGGAGTGATAGAAATC |
| Senescence | *Cdkn1a*$^{Var1}$ | AGATCCACAGCGATATCCAGACA | GTCGGACATCACCAGGATTGGAC |
| Senescence | *Cdkn1a*$^{Var2}$ | CACTTTGCCAGCAGAATAAAAGGTG | ACACTTTGCTCCTGTGCGGAAC |
| Senescence | *Tp53* | TCTTATCCGGGTGGAAGGAAA | GGCGAAAAGTCTGCCTGTCTT |
| ATTAC transgenic cassette | *FKBP-Casp8* | CAGATTATGCCTATGGTGCCACTG | GTAAACTTTGTCCAAAGTCTGTGATTCAC |
| Reference | *Actb* | AATCGTGCGTGACATCAAAGAG | GCCATCTCCTGCTCGAAGTC |
| Reference | *Gapdh* | GACCTGACCTGCCGTCTAGAAA | CCTGCTTCACCACCTTCTTGA |
| SASP | *Ager* | CTGGCACTTAGATGGGAAACT | TGTCTCCTGGTCTCTTCCTT |
| SASP | *Ccl2* | GTCTGTGCTGACCCCAAGAAG | TGGTTCCGATCCAGGTTTTTA |
| SASP | *Ccl3* | TCCCAGCCAGGTGTCATTTT | TTGGAGTCAGCGCAGATCTG |
| SASP | *Ccl5* | GCCCACGTCAAGGAGTATTTCT | ACAAACACGACTGCAAGATTGG |
| SASP | *Ccl7* | CCCTGGGAAGCTGTTATCTTCA | CTGATGGGCTTCAGCACAGA |
| SASP | *Ccl8* | CCACACAGAAGTGGGTCAGTGA | TTCAAGGCTGCAGAATTTGAGA |
| SASP | *Csf1* | ATTGCCAAGGAGGTGTCAGAA | GGACCTTCAGGTGTCCATTCC |
| SASP | *Csf2* | CCTTGAACATGACAGCCAGCTA | CACAGTCCGTTTCCGGAGTT |
| SASP | *Csf3* | CCTGCAGGCTCTATCGGGTAT | ATCCAGCTGAAGCAAGTCCAA |
| SASP | *Cxcl1* | CCGAAGTCATAGCCACACTCAA | CAAGGGAGCTTCAGGGTCAAG |
| SASP | *Cxcl10* | TGAATCCGGAATCTAAGACCA | TTTTTGGCTAAACGCTTTCAT |
| SASP | *Cxcl12* | GCCAACGTCAAGCATCTGAAA | CAGCCGTGCAACAATCTGAA |
| SASP | *Cxcl15* | TCCATGGGTGAAGGCTACTGT | TTCATTGCCGGTGGAAATTC |

| | | | |
|---|---|---|---|
| SASP | *Cxcl16* | GCACCCCTGCACATAGTCAGA | AGGACAGTGCTCCTGATGGAA |
| SASP | *Cxcl2* | TCAAGGGCGGTCAAAAAGTT | CAGTTAGCCTTGCCTTTGTTCA |
| SASP | *Fas* | CTGCACCCTGACCCAGAATAC | ACAGCCAGGAGAATCGCAGTA |
| SASP | *Fasl* | CCGCTCTGATCTCTGGAGTGA | CACGAAGTACAACCCAGTTTCG |
| SASP | *Foxo4* | CCACGAAGCAGTTCAAATGCT | TTCAGACTCCGGCCTCATTG |
| SASP | *Gas6* | AGGCTCAACTACACCCGAACA | TTTAACTTCCCAGGTGGTTTCC |
| SASP | *Gata4* | GCTCCTACTCCAGCCCCTAC | CAGGACTGGGCTGTCGAA |
| SASP | *Gdf15* | TGTGCAGGCAACTCTTGAAGA | GCGATACAGGTGGGGACACT |
| SASP | *Hmgb1* | TCCTTCGGCCTTCTTCTTGTT | AGGATGCTCGCCTTTGATTTT |
| SASP | *Icam1* | GTGGCGGGAAAGTTCCTGTT | GTCCAGCCGAGGACCATACA |
| SASP | *Ifng* | TTGGCTTTGCAGCTCTTCCT | ATGACTGTGCCGTGGCAGTA |
| SASP | *Igf1* | AAAAGCAGCCCGCTCTATCC | CTTCTGAGTCTTGGGCATGTCA |
| SASP | *Igfbp2* | GCCCCCTGGAACATCTCTACT | GTTGTACCGGCCATGCTTGT |
| SASP | *Igfbp3* | ACCTGCTCCAGGAAACATCAGT | TTTCCACACTCCCAGCATTG |
| SASP | *Igfbp4* | GCAACTTCCACCCCAAACAGT | CCTGTCTTCCGATCCACACA |
| SASP | *Il10* | TGGCTCAGCACTGCTATGCT | TGTACTGGCCCCTGCTGATC |
| SASP | *Il12a* | ATCCTGCTTCACGCCTTCAG | GATAGCCCATCACCCTGTTGA |
| SASP | *Il12b* | GCCAGTACACCTGCCACAAAG | TGTGGAGCAGCAGATGTGAGT |
| SASP | *Il15* | GGCATTCATGTCTTCATTTTGG | TCCAGTTGGCCTCTGTTTTAGG |
| SASP | *Il17a* | GGACTCTCCACCGCAATGAA | GCACTGAGCTTCCCAGATCAC |
| SASP | *Il1a* | AAGAGACCATCCAACCCAGATC | CCTGACGAGCTTCATCAGTTTG |
| SASP | *Il1b* | TCAGGCAGGCAGTATCACTCA | CACGGGAAAGACACAGGTAGCT |
| SASP | *Il3* | GCCTGCCTACATCTGCGAAT | CGAAAGTCATCCAGATCTCGAA |
| SASP | *Il4* | TCCTCACAGCAACGAAGAACAC | AAGCACCTTGGAAGCCCTACA |
| SASP | *Il6* | ACCACGGCCTTCCCTACTTC | TTGGGAGTGGTATCCTCTGTGA |

| | | | |
|---|---|---|---|
| SASP | *Il8* | TCCATGGGTGAAGGCTACTGT | TTCATTGCCGGTGGAAATTC |
| SASP | *Inhba* | CAGGAAGACACTGCACTTTGA | TTCAGGAAGAGCCACACTTCT |
| SASP | *Irf1* | CAGCCGAGACACTAAGAGCAAA | GAGAAAGTGTCCGGGCTAACAT |
| SASP | *Lif* | GCTGTATCGGATGGTCGCATA | TCTGGTCCCGGGTGATATTG |
| SASP | *Mif* | GCCACCATGCCTATGTTCATC | GGGTGAGCTCCGACAGAAAC |
| SASP | *Mmp12* | GTGCCCGATGTACAGCATCTT | GGTACCGCTTCATCCATCTTG |
| SASP | *Mmp13* | TGAGGAAGACCTTGTGTTTGCA | GCAAGAGTCGCAGGATGGTAGT |
| SASP | *Mmp2* | TGTGGGTGGAAATTCAGAAGGT | ACTTGTTGCCCAGGAAAGTGA |
| SASP | *Mmp7* | TTGCTGCCACCCATGAATTT | TCACAGTACCGGGAACAGAAGA |
| SASP | *Mmp8* | TGGCTGCTCATGAATTTGGA | CATCAAGGCACCAGGATCAGT |
| SASP | *Mmp9* | TGAGTCCGGCAGACAATCCT | CCCTGGATCTCAGCAATAGCA |
| SASP | *Nfkb1* | GGCTTTGCAAACCTGGGAAT | TCCGTGCTTCCAGTGTTTCA |
| SASP | *Pappa* | CATCTCAGGTGTGTCGAACCA | TGCAAGGATACCAAGCATGCT |
| SASP | *Pdgfa* | CTCGAAGTCAGATCCACAGCAT | CAGCCCCTACGGAGTCTATCTC |
| SASP | *Pdgfb* | GCTGAGCTGGACTTGAACATGA | CCTCGAGATGAGCTTTCCAACT |
| SASP | *Serpinb2* | TTCCGCATACTGGAAACATCAG | GGATGCGTCCTCAATCTCATC |
| SASP | *Serpine1* | GGACACCCTCAGCATGTTCA | CGGAGAGGTGCACATCTTTCT |
| SASP | *Sparc* | GAGGAGGTGGTGGCTGACAA | CACCTTGCCATGTTTGCAAT |
| SASP | *Spp1* | GAGGAGGTGGTGGCTGACAA | CACCTTGCCATGTTTGCAAT |
| SASP | *Tgfb1* | AGCGCTCACTGCTCTTGTGA | GCTGATCCCGTTGATTTCCA |
| SASP | *Tgfb2* | GGTGGCGCTCAGTCTGTCTAC | TCTTGCGCATAAACTGATCCAT |
| SASP | *Tgfb3* | CTGTCCACTTGCACCACGTT | CCTAATGGCTTCCACCCTCTT |
| SASP | *Tgfbr1* | CGTGTGCCAAATGAAGAGGAT | AAGGTGGTGCCCTCTGAAATG |
| SASP | *Tnf* | GTTCTGCAAAGGGAGAGTGG | GCACCTCAGGGAAGAGTCTG |
| SASP | *Tnfrsf11b* | CCAAGAGCCCAGTGTTTCTT | CCAAGCCAGCCATTGTTAAT |

| | | | |
|---|---|---|---|
| SASP | *Tnfrsf12a* | CCGCCGGAGAGAAAAGTTTAC | GGGTGCTCCTCACTGGATCA |
| SASP | *Tnfsf10* | CTCTCGGAAAGGGCATTCATT | TCGATGACCAGCTCTCCATTC |
| SASP | *Tnfsf11* | GCTGGGACCTGCAAATAAGT | TTGCACAGAAAACATTACACCTG |
| SASP | *Vcam1* | GGCTCCAGACATTTACCCAGTT | CATGAGCTGGTCACCCTTGAA |
| SASP | *Vegfa* | GTACCTCCACCATGCCAAGTG | TGGGACTTCTGCTCTCCTTCTG |

**Supplementary Table 3.** Assessment of sex as a biological variable. 2-way ANOVA results found no interaction effects of sex on primary endpoints of fracture healing.

| Metric | Source of Variation | % of total variation | P value | P value summary | Significant? |
|---|---|---|---|---|---|
| Wehrle score | Interaction | 0.1703 | 0.5303 | ns | No |
| | Sex | 0.1482 | 0.0597 | ns | No |
| | Timepoint | 91.34 | <0.0001 | **** | Yes |
| Relative callus area | Interaction | 0.3442 | 0.9887 | ns | No |
| | Sex | 0.3583 | 0.4390 | ns | No |
| | Timepoint | 26.03 | <0.0001 | **** | Yes |
| qPCR *Cdkn1a* | Interaction | 1.181 | 0.6265 | ns | No |
| | Sex | 0.3526 | 0.7895 | ns | No |
| | AP Treatment | 46.51 | 0.0094 | ** | Yes |

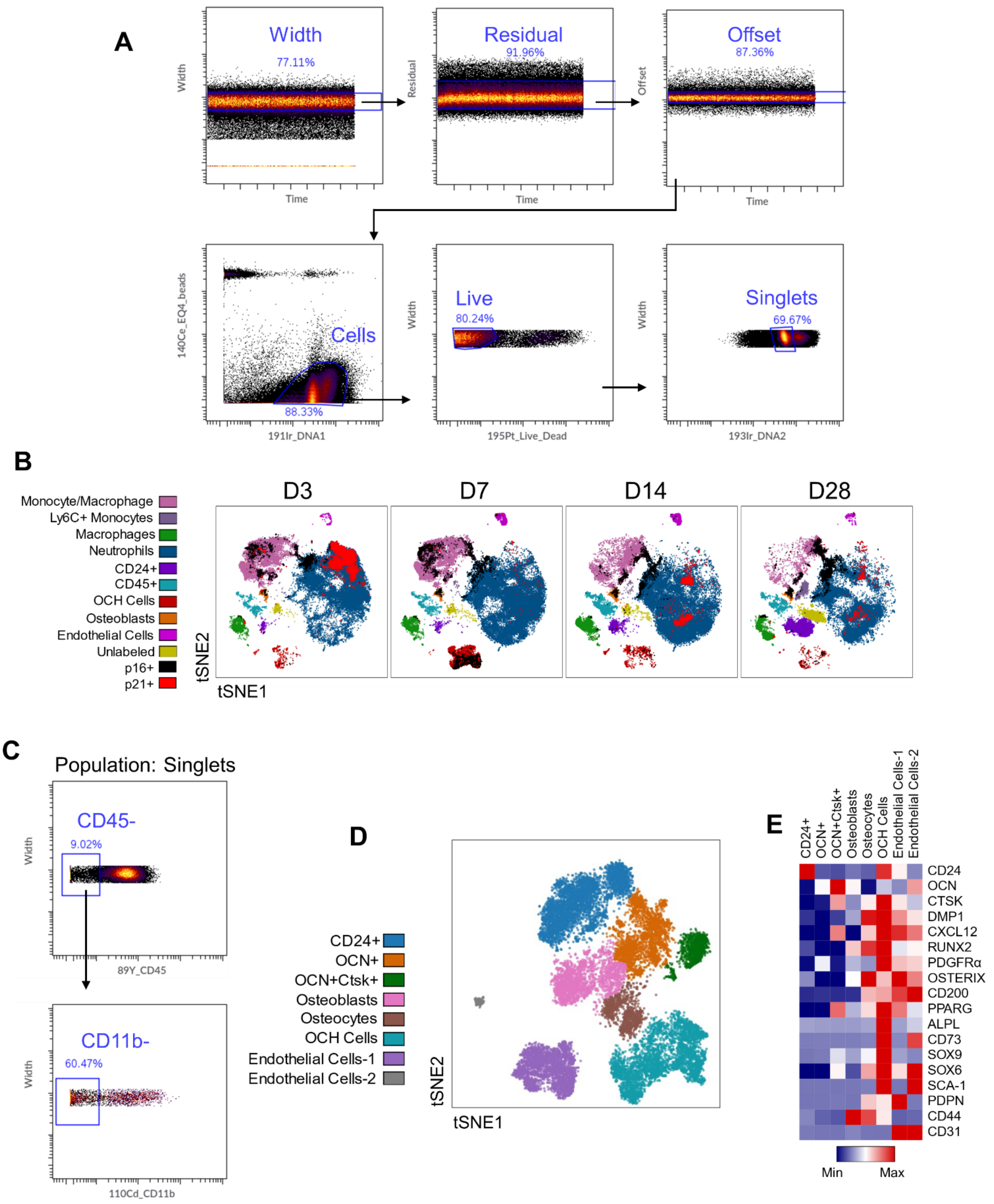


**Supplementary Figure 1: CyTOF characterization of murine fracture healing.** (A) CyTOF gating strategy for purification of live singlets. (B) tSNE visualization of clustered fracture callus cell populations by CyTOF, overlaid with p16+ (black) or p21+ (red) cells. (C) Gating strategy for purification of CD45-CD11b- cells. (D) tSNE visualization of re-clustered CD45-CD11b- non-immune callus cells. (E) Heatmap of identity marker mean expression in each cluster. Day 3: n=12 mice (6 female, 6 male), day 7: n=12 mice (6 female, 6 male), day 14: n=12 mice (6 female, 6 male), day 28: n=11 mice (6 female, 5 male).

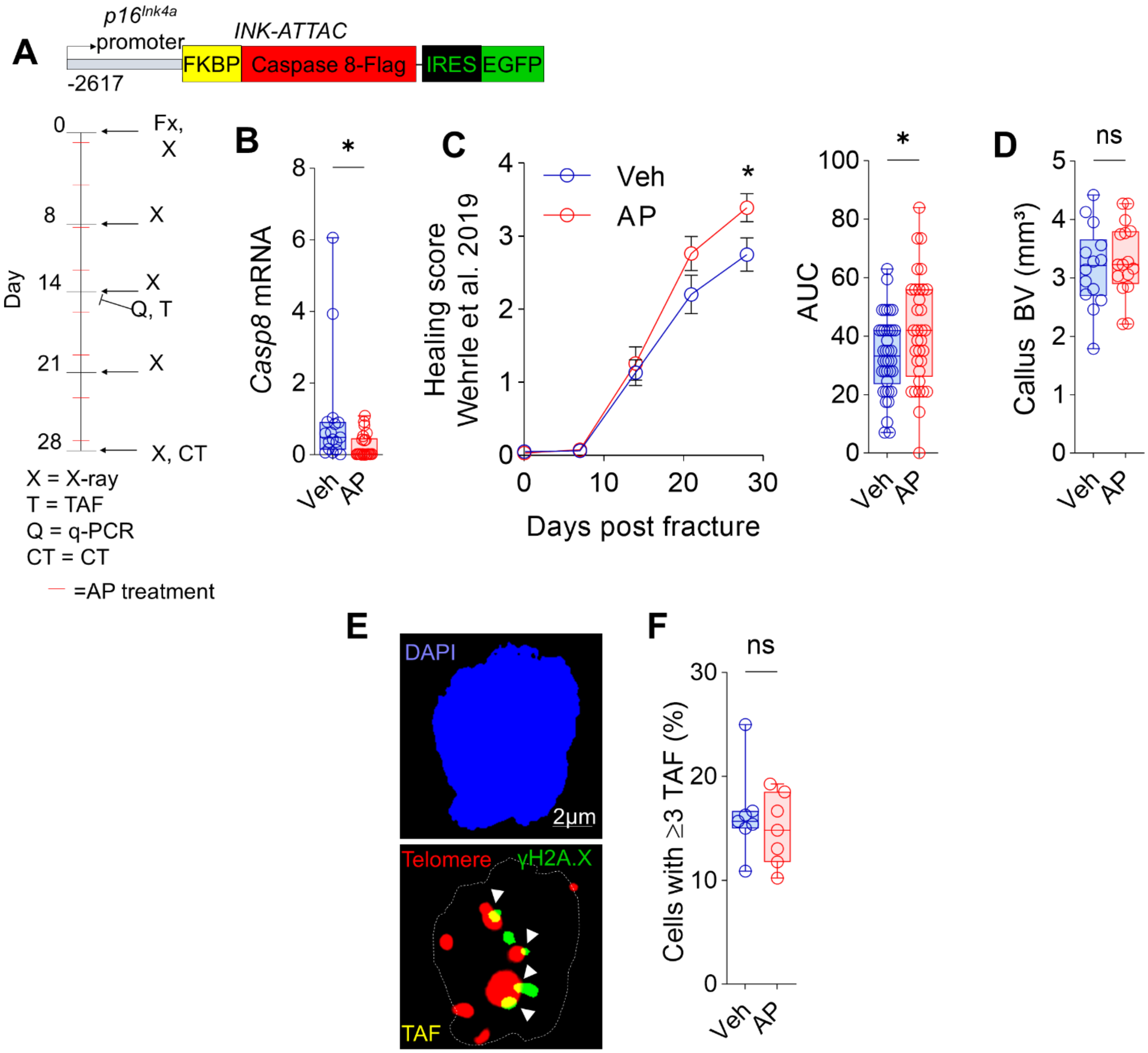


**Supplementary Figure 2: p16+ cell clearance minimally affects murine fracture healing.** (A) Schematic of the transgene in the *p16-INK-ATTAC* mice and overall study design. (B) qPCR measurement of INK-ATTAC transgene (*Casp8*), reduced with AP treatment (C) X-ray healing score and area under the curve (AUC) analysis in Vehicle- and AP-treated *p16-INK-ATTAC* mice over a 5-week healing period. (D) Micro-CT analysis of callus bone volume. (E) TAF (yellow, see arrowheads) were defined as sites of γH2A.X-associated DNA damage co-localized with telomeres (n=70 cells were analyzed per bone). (F) Quantification of cells with ≥3 TAFs. (B) n=17 Veh (9 female, 8 male), n=22 AP (12 female, 10 male). (C) n=40 Veh (22 female, 18 male), n=33 AP (20 female, 13 male). (D) n=14 Veh (10 female, 4 male), n=16 AP (9 female, 7 male). (E-F) n=7 Veh (5 female, 2 male), n=7 AP (5 female, 2 male). (B, D, F) Mann-Whitney or unpaired t test as appropriate. (C) Two-way ANOVA with Sidak multiple comparisons test. *$p<0.05$.

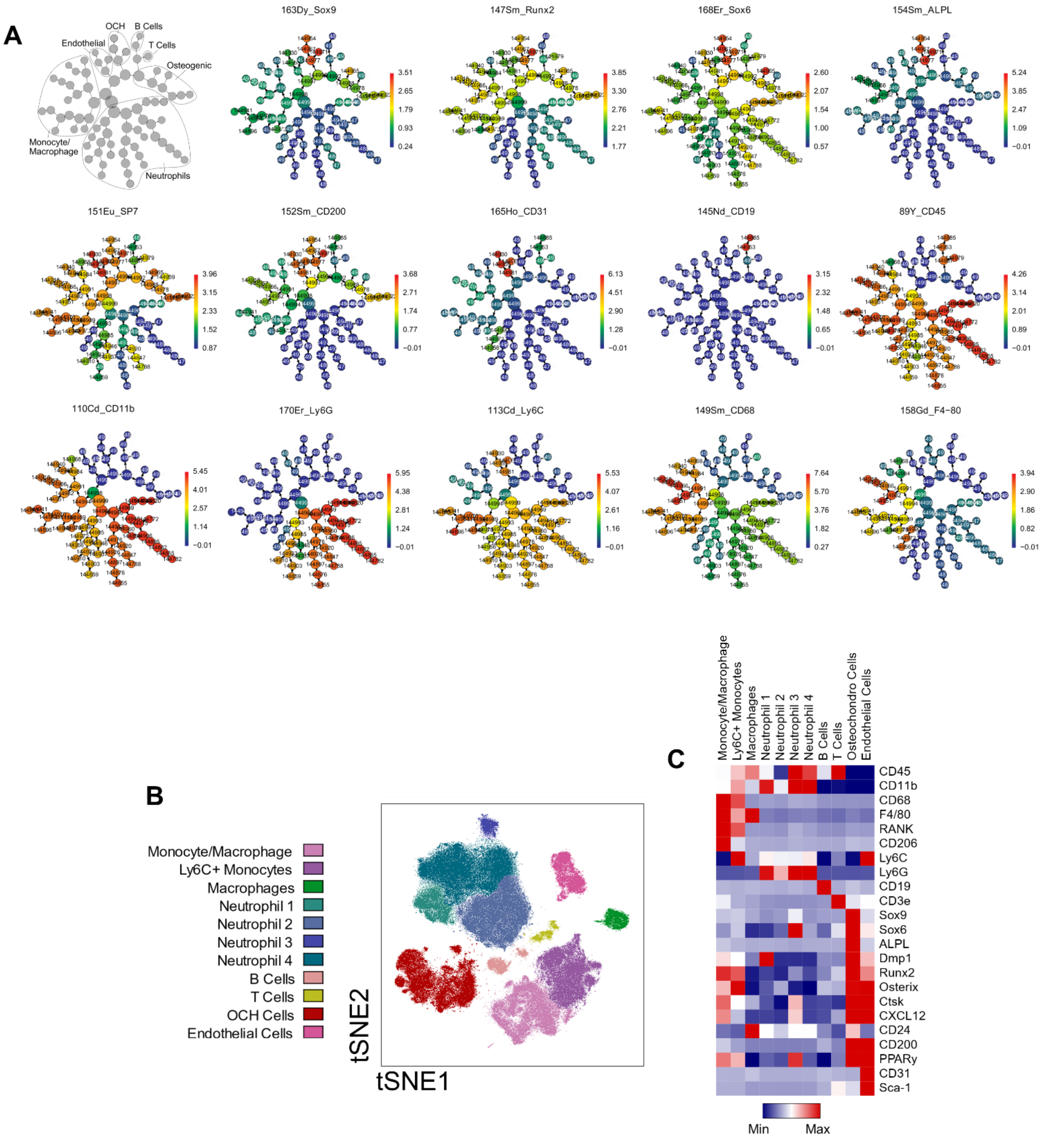


**Supplementary Figure 3: CyTOF characterization of fracture callus cells in *p21-ATTAC* mice.** (A) CITRUS expression plots of identity markers. (B) tSNE visualization of all clustered callus cell populations merged from vehicle- or AP-treated *p21-ATTAC* mice at D7 post-fracture. (C) Heatmap of identity marker mean expression in each cluster. n=16 Veh (8 female, 8 male), n=13 AP (7 female, 6 male).

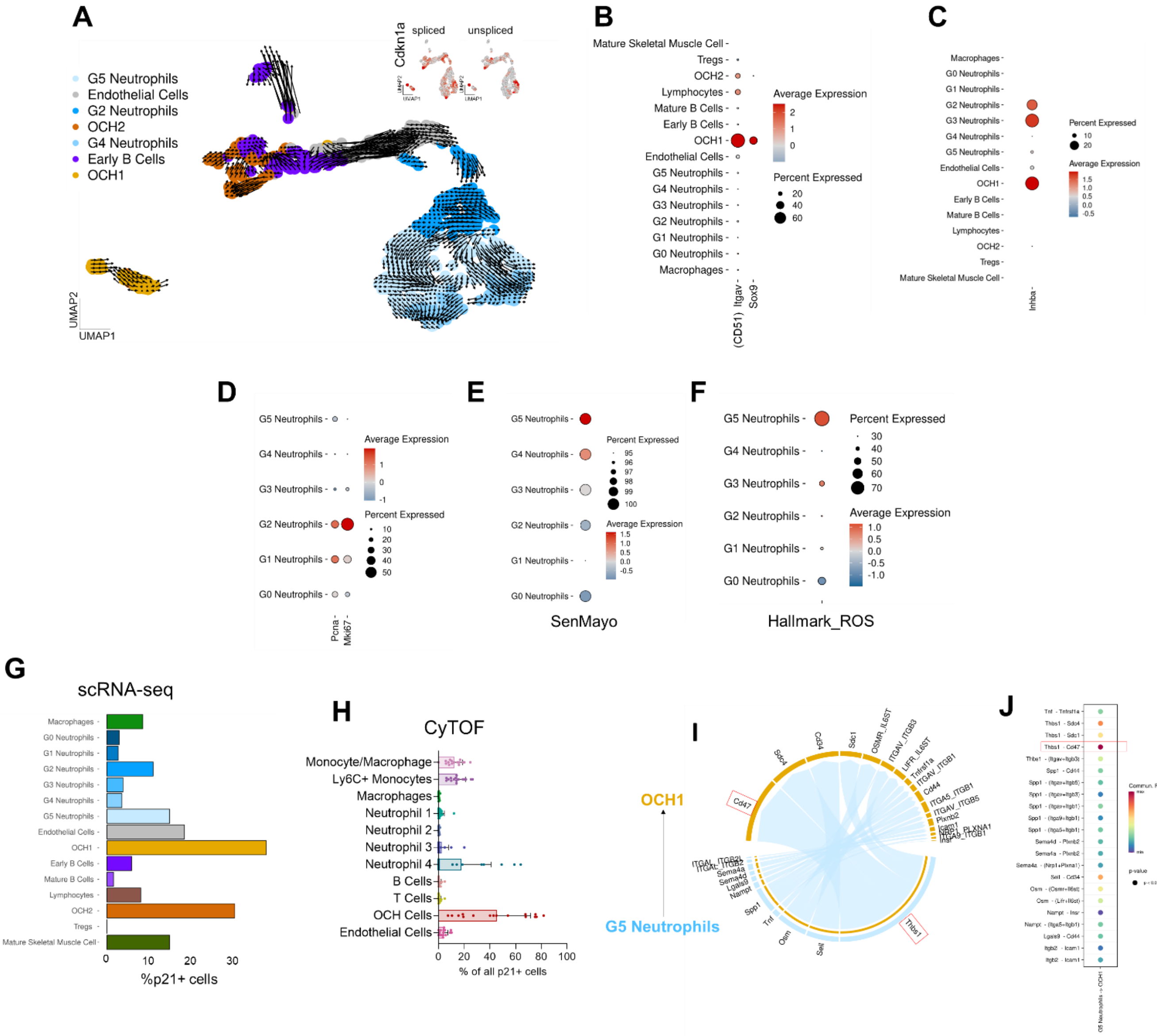


**Supplementary Figure 4: Detailed analysis of callus cell scRNA-seq.** (A) RNA-Velocity[68] demonstrates the detached state of OCH1 cells (lower left), while neutrophil development is gradually shown on the right developmental path. (B) OCH1 cells are highly enriched in *Itgav* and *Sox9*, while (C) high in *Inhba*, as are G2 and G2 neutrophils. (D) Proliferation markers (*Pcna* and *Mki67*) are highest in the G2 neutrophil population, but not in the G5 population. (E) The neutrophil population with the highest SenMayo score is G5, which also (F) enriches the most in the Hallmark reactive oxygen species (ROS, M5938). (G) In the scRNA-Seq dataset, the percentage of p21+ cells within the neutrophil clusters is highest in G5, which is (H) mirrored in the CyTOF dataset, where the Neutrophil 4 cells show the highest percentage of p21+ cells. (I) The communicational path preferably used by G5 neutrophils to OCH1 cells is the *Thbs1*/*Cd47* pathway, that is (J) highly enriched for these two cell types.

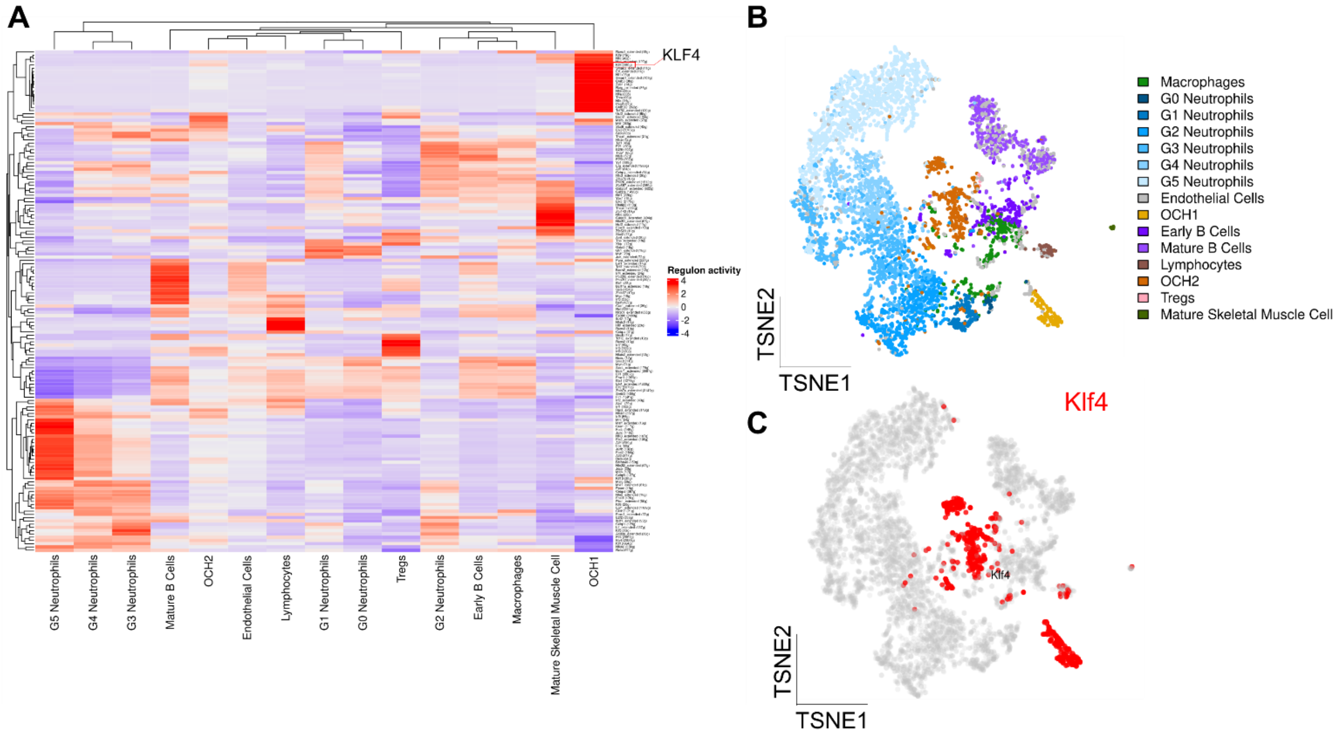


**Supplementary Figure 5: SCENIC regulatory analysis of callus cell populations.** (A) All significantly regulating regulons are depicted for each cell type. Notably, *Klf4* is among the key controlling genes for the OCH1 and OCH2 cells. (B) tSNE visualization after regulon assessment. (C) The *Klf4* regulon alone controls large parts of the OCH1 and OCH2 clusters.

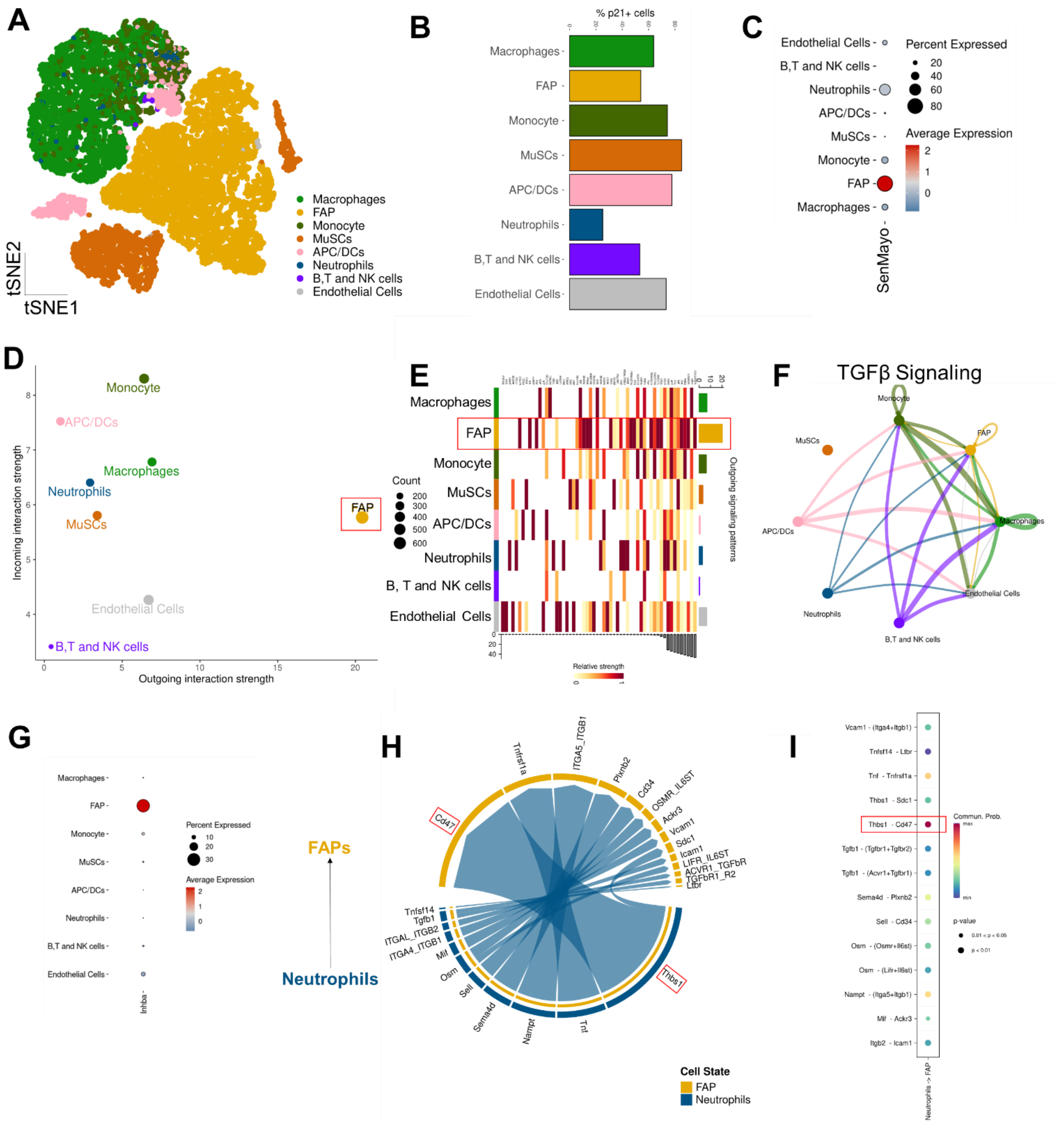


**Supplementary Figure 6: Detailed analysis of skeletal muscle scRNA-seq** (A) tSNE plot demonstrating the heterogeneity of murine skeletal muscle (GSE197017).[12] (B) The percentage of p21+ cells is highest in muscle stem cells (MuSCs) and antigen-presenting/dendritic cels (APC/DCs), while (C) SenMayo is mostly enriched in the fibroadipogenic progenitors (FAPs). (D, E) These FAPs are the cells with the highest outgoing interaction strength. (F) Amongst others, the *TGFβ* pathway seems to be crucial in the FAPs. (G) The FAPs are also highest in *Inhba*. (H) Similar to bone, neutrophils primarily communicate to these FAPs via the *Thbs1*/*Cd47* pathway that is (I) highly enriched for the communication neutrophils – FAPs.
